# Misrouting of Glucagon and Stathmin-2 Towards Lysosomal System of α-Cells in Glucagon Hypersecretion of Diabetes

**DOI:** 10.1101/2021.04.08.439083

**Authors:** Farzad Asadi, Savita Dhanvantari

**Affiliations:** Department of Pathology and Laboratory Medicine, Schulich School of Medicine & Dentistry, Western University, London, ON, Canada; Department of Medical Biophysics, Western University, London, ON, Canada; Lawson Health Research Institute, London, ON, Canada

**Keywords:** Glucagon secretion, Hyperglucagonemia, Diabetes, Stathmin-2, Lysosome, Integrin

## Abstract

Glucagon hypersecretion from the pancreatic α-cell is a characteristic sign of diabetes, which exacerbates fasting hyperglycemia. Thus, targeting glucagon secretion from α-cells may be a promising approach for combating hyperglucagonemia. We have recently identified stathmin-2 as a protein that resides in α-cell secretory granules, and showed that it regulates glucagon secretion by directing glucagon towards the endolysosomal system in αTC1-6 cells. Here, we hypothesized that disruption of Stmn2-mediated trafficking of glucagon to the endolysosomes contributes to hyperglucagonemia. In isolated islets from male mice treated with streptozotocin (STZ) to induce diabetes, Arg-stimulated secretion of glucagon and Stmn2 was augmented. However, cell glucagon content was significantly increased (p<0.001), but Stmn2 levels were reduced (p<0.01) in STZ-treated mice, as measured by both ELISA and immunofluorescence intensity. Expression of *Gcg* mRNA increased ~4.5 times, while *Stmn2* mRNA levels did not change. Using confocal immunofluorescence microscopy, the colocalization of glucagon and Stmn2 in Lamp2A^+^ lysosomes was dramatically reduced (p<0.001) in islets from diabetic mice, and the colocalization of Stmn2, but not glucagon, with the late endosome marker, Rab7, significantly (p<0.01) increased. Further studies were conducted in αTC1-6 cells cultured in media containing high glucose (16.7 mM) for two weeks to mimic glucagon hypersecretion of diabetes. Surprisingly, treatment of αTC1-6 cells with the lysosomal inhibitor bafilomycin A1 reduced K^+^-induced glucagon secretion, suggesting that high glucose may induce glucagon secretion from another lysosomal compartment. Both glucagon and Stmn2 co-localized with Lamp1, which marks secretory lysosomes, in cells cultured in high glucose. We propose that, in addition to enhanced trafficking and secretion through the regulated secretory pathway, the hyperglucagonemia of diabetes may also be due to re-routing of glucagon from the degradative Lamp2A^+^ lysosome towards the secretory Lamp1^+^lysosome.

## Introduction

In diabetes, glucagon secretion from the pancreatic α-cell becomes abnormally up-regulated, resulting in hyperglucagonemia. This paradoxical glucagon hypersecretion from α-cells then exacerbates hyperglycemia (Honzawa et al., 2019; Salehi et al., 2006; Unger and Cherrington, 2012). It has been suggested that, in order to fully control hyperglycemia in diabetes, glucagon secretion should be suppressed. Therefore, the mechanisms and pathways that underlie the abnormal secretion of glucagon must be elucidated, so that potential targets for suppressive therapy can be identified.

Studies have shown that glucagon secretion can be controlled by targeting mediators of intracellular signaling and exocytosis within the α-cell. Agonists of the glucagon-like peptide 1 receptor (GLP-1R) inhibit glucagon secretion by directly acting on α-cells, or indirectly through releasing insulin, Zn^2+^, and GABA from β-cells, or by releasing somatostatin from δ-cells (DJ Drucker, 2018). GABAA receptor agonists also directly inhibit glucagon secretion through binding with GABAA receptor on α-cells, increasing Cl^-^ influx, and hyperpolarization of the plasma membrane (Feng et al., 2017; Rorsman et al., 1989). Antagonists of the glucosedependent insulinotropic peptide receptor (GIP-R) directly suppress the glucagonotropic effect of GIP on α-cells (Gasbjerg et al., 2018). Insulin directly suppresses glucagon secretion through binding its receptor on α-cells (Kawamori and Kulkarni, 2009; Kawamori et al., 2009) or indirectly by increasing secretion of somatostatin from δ-cells (Vergari et al., 2019). Blockers of K_ATP_ channels inactivate ion channels, and reduce α-cell electrical activity, which result in suppression of glucagon secretion (MacDonald et al., 2007; Olsen et al., 2005; Zhang et al., 2013). As well, in addition to blocking the effects of glucagon on hepatic glucose mobilization, anti-glucagon receptor antibodies may also block the autocrine effect of glucagon on glucagon secretion from α-cells (Lee et al., 2016; Wang et al., 2015).

All of the above mechanisms eventually converge on the α-cell secretory pathway by which glucagon is stored in and secreted from secretory granules. We hypothesize that elucidating the intracellular mechanisms of glucagon trafficking through the α-cell secretory pathway could also yield clues on possible mechanisms of the regulation of glucagon secretion. Using αTC1-6 cells, we have shown that components of the regulated secretory pathway are up-regulated after chronic exposure to high levels of glucose (McGirr et al., 2005) and that proteins that associate with glucagon within secretory granules (glucagon interactome) play a role in the regulation of glucagon secretion (Asadi and Dhanvantari, 2019). We showed that the interactome was responsive to glucose, GABA and insulin, known modulators of glucagon secretion. In particular, treatment of αTC1-6 cells with insulin, which can override glucose-mediated regulation of glucagon secretion (Kawamori et al., 2009), appeared to recruit stathmin-2 (Stmn2 or SCG10) to the interactome, thus potentially identifying another inhibitor of glucagon secretion. In a subsequent study, we showed that Stmn2 is co-secreted in a regulated manner with glucagon, and when over-expressed, suppressed glucagon secretion by increasing its trafficking through the endolysosomal pathway in αTC1-6-cells (Asadi and Dhanvantari, 2020).

There is one report on the reuptake of glucagon by α-cells after secretion and trafficking to the endolysosomal system for degradation (Amherdt et al., 1989). Our previous study was the first to show that the endolysosomal system may figure prominently in the intracellular trafficking of glucagon as it is directed through the regulated secretory pathway(Asadi and Dhanvantari, 2020). Interestingly, parallel findings in β-cells showed that there is a shuttling of proinsulin from secretory pathway towards the endolysosomal system for degradation, which may be an underlying mechanism for β-cell failure in type 2 diabetes (Pasquier et al., 2019). Therefore, the endolysosomal trafficking may be a novel pathway for the dysregulation of both insulin and glucagon secretion in diabetes. We hypothesize that a disruption in the trafficking of glucagon and Stmn2 through the endolysosomal pathway might be a possible mechanism by which glucagon secretion becomes dysregulated in diabetes, resulting in glucagon hypersecretion and hyperglucagonemia.

In the present study, we examined the endolysosomal trafficking of glucagon and Stmn2 in islets from streptozotocin (STZ)-treated male mice, and in αTC1-6 cells cultured in high glucose conditions. We report that, in STZ-treated mice, a relative reduction of Stmn2 is associated with inhibition of the trafficking of glucagon from the late endosome to degradative Lamp2A^+^ lysosomes. Further studies in αTC1-6 cells cultured for a long-term in media containing high glucose levels suggest that glucagon is re-routed to a secretory lysosomal subpopulation expressing Lamp1. Therefore, we propose that alterations in the endolysosomal trafficking of glucagon contributes to the hyperglucagonemia of diabetes.

## Materials and Methods

### Animals

C57BL/6 male mice (8 weeks old) were purchased from the Jackson Laboratory (Bar Harbor, Maine, United States) and housed in the Lawson Health Research Institute Animal House Facility (London, ON, Canada). Mice were kept at 12h light/12h dark cycle and had access to water and regular chow diet ad libitum. All mice were treated and euthanized in accordance with the guidelines set out by the Animal Use Subcommittee of the Canadian Council on Animal Care at Western University based on the approved Animal Use Protocol AUP 2012-020. Mice were fasted 5h before blood collection or euthanasia.

### Induction of diabetes

Streptozotocin (STZ; Cat# S0130, Sigma) was dissolved in freshly prepared 0.1 M sodium citrate buffer, pH 4.5, and immediately used. Mice (n=18) were intraperitoneally injected with 30 mg/kg STZ for 5 consecutive days and used for microscopy (total n=7; 7 out of 7 for immunofluorescent microscopy; 4 out of 7 for transmission electron microscopy), islet secretion (n=7) and gene expression (n=4) studies. Three days after the last STZ injection, blood was sampled by tail vein lancing and glucose levels were determined with the OneTouch Ultra glucometer. Values above 14 mmol/L were considered an indicator of diabetes onset (https://www.jax.org). Control mice (n=18) were injected with citrate buffer alone at the same regimen. Animals were euthanized 14 days after the last STZ injection by cervical dislocation under deep isoflurane anesthesia.

### Blood collection

Prior to cervical dislocation and under isoflurane induced anesthesia, blood (1 mL) was collected by cardiac puncture into microcentrifuge tubes containing 15 μL of anticoagulant (15% Na_2_-EDTA) and 15 μL of freshly prepared enzyme inhibitor cocktail (Cat# 4693159001 Millipore Sigma). Samples were kept on ice and then centrifuged at 1500×g for 15 min at 4°C. Plasma was collected and kept at −80°C until analysis.

### Preparation of pancreas tissue sections for confocal immunofluorescence microscopy

Immediately after euthanasia, pancreata of STZ-treated (n=7) and vehicle-treated (n=7) mice were excised, fixed in 10% buffered formalin for 3 days and treated with 70% ethanol for one day before paraffin embedding. Paraffin-embedded tissue blocks were longitudinally sectioned in 5 μm slices and fixed on glass microscope slides. The tissue samples were de-paraffinized by graded washes using xylene, ethanol and PBS. Antigen retrieval was conducted in sodium citrate buffer (10 mM sodium citrate, 0.05% Tween 20, pH 6) with 20 min steam heating of slides in a steam cooker. After permeabilization with 0.1% Triton X-100 in PBS, Background Sniper (Cat# BS966H, Biocare Medical) was used to block non-specific background staining. Samples were incubated with primary antibodies against glucagon (Cat # ab10988 or Cat# ab92517, Abcam; 1:1000), Stmn2 (Cat # ab115513, Abcam; 1:250 or Cat# 720178, Thermo Fisher Scientific; 1:500), insulin (Cat# ab7842, Abcam; 1:500 or Cat# I2018, Sigma; 1:1000), late endosome marker, Rab7 (Cat# ab126712, Abcam; 1:500), lysosomal marker, Lamp2A (Cat# ab18528, Abcam; 1:1000), recycling endosome markers Rab11A (Cat# ab180778, Abcam; 1:100) and Rab11B (Cat# ab228954, Abcam;1:100). Then, corresponding secondary antibodies used were donkey anti-goat IgG Alexa Fluor 555 (Cat# ab150130, Abcam; 1:500), donkey anti-rabbit 488 (Cat#ab150073, Abcam; 1:500), and donkey anti-mouse 555 (Cat# A-31570, Molecular Probes; 1:500). Nucleus counter-staining was done by DAPI and coverslips were mounted using Prolong Antifade mountant (Cat# P36982, Thermo Fisher Scientific). As a background control for Stmn2, pancreas sections were incubated with only the corresponding secondary antibody.

### Image acquisition

Images were acquired through Nikon A1R Confocal microscope with a ×60 NA plan-Apochromat oil differential interference contrast objective and NIS-Elements software (Nikon, Mississauga, Canada). Acquisition of high-resolution images was done by selecting Nyquist XY scan area, 1024×1024-pixel size scanning of the selected area and 2DDeconvolution of the captured images, as we have done previously(Asadi and Dhanvantari, 2020).

### Image Analysis

Three adjacent longitudinal slices of pancreas were placed on each glass slide. In total, 10 slides were prepared from each pancreas. Image analysis was performed by NIS-Elements software (Nikon, Mississauga, Canada). To calculate colocalization values of endosomal and lysosomal markers with glucagon or Stmn2 within the same islet, channels were pseudocolored for Lamp2A, Rab7, Rab11A or Rab11B. Colocalization of pixels from each pseudocolored image was used to calculate Pearson’s correlation coefficient (PCC), as we have done previously (Asadi and Dhanvantari, 2019; Guizzetti et al., 2014). Regions of interest (ROIs) were manually drawn around each islet and then defined PCC values for colocalization between Stmn2 and target markers (glucagon, insulin, Lamp2A, Rab7, Rab11A, Rab11B) were calculated using the colocalization algorithm of NIS-Elements software. To show the relationships between expression levels of Stmn2 and the target markers (glucagon, insulin, Lamp2A, Rab7, Rab11A, Rab11B) in α or β-cells of the pancreatic islet, binary images were generated using M-Threshold algorithm of NIS-Elements software. ROIs were manually drawn around each binary arranged image of the islet and the fluorescence intensity of each marker was calculated. Levels of fluorescence intensities were normalized by dividing by the intensity of DAPI within each ROI. These values were used for linear regression analysis between Stmn2 and the target markers or t-test analysis between non-diabetic and diabetic α-cells.

### Double immunogold labeling transmission electron microscopy

Double immunogold labeling TEM was done based on the protocol of Aida et al (2014) with some modifications (Aida et al., 2014) as we have recently used (Asadi and Dhanvantari, 2020). Briefly, in both nondiabetic (n=4) and diabetic (n=4) mice, pancreas was dissected, and a piece of pancreas in its long axis was cut, and immediately placed into McDowell Trump’s fixative (Cat# 18030-10, Electron Microscopy Sciences) for 1h. Then, after washing with PBS, samples were cut into smaller pieces, and dehydrated in the increasing concentrations of ethanol (10%, 20%, 30%, 50%, 70%, 90%, 100% and 100%) at 30 min per concentration. Samples were sequentially embedded in LR White Resin (Cat# 14381, Electron Microscopy Sciences) as follows: ethanol-LR White mixture A (3:1, v/v; 2h), ethanol-LR White mixture B (1:1, v/v; 8h), ethanol-LR White mixture C (1:3, v/v; 12h), and 3 × 12h in pure LR White. Samples were then placed into a beem capsule, filled with pure LR White and incubated at 50°C for 24h. Semi-thin sections (500 nm) were cut from embedded samples for Toluidin blue staining (1% Toluidin blue for 2 min). After defining the position of the islet within the pancreatic tissue, ultra-thin section slices (70 nm) were prepared using a diamond microtome. The sections were mounted on formvar-carbon coated nickel grid (300 meshes; Cat# FCF300-NI, Electron Microscopy Sciences). Afterwards, slices were washed with Tris-buffered saline containing 0.1% Tween 20 (TBS-T) and incubated in blocking buffer (2% BSA in PBS plus 0.05% Tween 20) for 30 min at room temperature. Slices were incubated with primary antibodies (1:10 in blocking buffer) against glucagon (Cat# ab10988; Abcam) and Stmn2 (Cat# ab115513; Abcam) at 4°C overnight. After washing with TBS-T, slices were incubated with gold-conjugated secondary antibodies of donkey anti-goat (18 nm; cat# ab105270, Abcam; 1:50) and donkey anti-mouse (10nm; cat# ab39593, Abcam; 1:50) for 2h at room temperature. After washing with TBS-T and staining with Uranyless (Cat# 22409, Electron Microscopy Sciences), TEM imaging was conducted at the Biotron Experimental Research Center, Western University, London, ON, Canada.

### Proglucagon and stathmin-2 gene expression

Handpicked islets (~180) from control (n=4) and diabetic (n=4) mice were placed into 1 mL Trizol (Cat# 15596018, Ambion) and processed for RNA extraction, as described previously (Augereau et al., 2016)(Li et al., 2014). Islets were homogenized by being passed 10 times through a 25-gauge needle, and again through a 27-gauge needle. After centrifugation at 10000×g for 5 min at 4°C, the supernatant was mixed with chloroform, vortexed for 30 seconds and placed on ice for 2 min. After centrifugation at 12000×g for 15 min at 4°C, the aqueous layer was collected and mixed with 0.5 volumes of high salt solution (0.8 M Na-citrate containing 1.2 M NaCl). Isopropanol (0.5 volumes) was added, and the samples were mixed, incubated for 10 min at room temperature and centrifuged at 12000×g for 30 min at 4°C. The pellet was dissolved in 70% ethanol and RNA was purified by RNeasy kit (Cat # 74104, Qiagen) according to the supplier’s protocol. cDNA synthesis was performed using the SuperScript III First Strand Synthesis Supermix for qRT-PCR (Cat # 11752050, Thermo Fisher Scientific), according to the manufacturer’s protocol. Real-time PCR was performed using Quant Studio Design and Analysis Real-Time PCR Detection System in conjunction with the Maxima SYBR Green qPCR Master Mix (Cat # K0221, Thermo Fisher Scientific) using specific primers for *Stmn2*: forward, 5’-GCAATGGCCTACAAGGAAAA-3’; reverse, 5’-GGTGGCTTCAAGATCAGCTC-3’; *Gcg*: forward, 5’-AACAACATTGCCAAACGTCA-3’; reverse, 5’-TGGTGCTCATCTCGTCAGAG-3’ and *18S rRNA*: forward, 5’-ACGATGCCGACTGGCGATGC-3’; reverse, 5’-CCCACTCCTGGTGGTGCCCT-3’. Gene expression levels were normalized to that of *18S rRNA*. Gene expression in the diabetic condition was normalized to the corresponding control group and expressed as percent of matched control. Statistical analysis was performed using t-test at α = 0.05.

### Primary islet culture

Islets were isolated and cultured as we have done previously (20) using a modified protocol from Li et al. (Li et al., 2009). Non-diabetic (n=7) or diabetic mice (n=7) were euthanized. The abdominal cavity was opened and 3 mL of 1.87 mg/mL collagenase V (Cat# C9263, Sigma;) in Hanks’ Balanced Salt Solution (HBSS) was injected into the common bile duct. The pancreas was then removed, placed into a Falcon tube containing 2 mL of the ice-cold collagenase V solution and incubated for 12 min at 37°C with occasional shaking. Digestion was stopped by adding 1 mM CaCl_2_ and the cell suspension was washed twice in 1 mM CaCl_2_. Islets were collected into a sterile petri dish using a 70 μm cell strainer with RPMI1640 containing 11 mM glucose plus 20 mM glutamine, 10% FBS and penicillin (110U/mL) and streptomycin (100 μg/mL). A total of 180 islets were handpicked under a stereomicroscope and incubated for 2h at 37°C. The medium was then changed to RPMI1640 containing 11 mM glucose plus 10% FBS and penicillin (110U/mL) and streptomycin (100 μg/mL) and islets were cultured overnight at 37°C.

### Islet secretion experiments

Glucagon and Stmn2 secretion from islets was measured based on the protocol by Suckow et al (2014). Briefly, islets were washed three times using Krebs-Ringer bicarbonate (KRB) buffer (135 mM NaCl, 3.6 mM KCl, 5 mM NaHCO_3_, 0.5 mM NaH_2_PO_4_, 0.5 mM MgCl_2_, 1.5 mM CaCl_2_, 10 mM HEPES; pH 7.4) containing 11 mM glucose, and then preincubated in this KRB for 1h. Islets were then incubated in KRB containing 1mM glucose in the presence or absence of arginine (25 mM) for 20 min. Media were collected into microcentrifuge tubes containing enzyme inhibitors (PMSF, 45 mM; Aprotinin, 5 μg/mL and sodium orthovanadate, 1mM). Samples were centrifuged at 14000 × g for 5 min at 4°C and the supernatant was collected and kept at −80°C until analysis. Islets were lysed in lysis buffer (0.1 M citric acid, 1% Triton X-100 plus enzyme inhibitors) and homogenized by passing 10 times through a 25-gauge needle, and again through a 27-gauge needle. The extracts were centrifuged at 14000×g for 15 min at 4°C and the supernatant was collected and kept at −80°C until analysis (Pisania et al., 2010). Glucagon levels in the media and islet extracts were determined by ELISA (Cat # 81520, Crystal Chem) according to the manufacturer’s instructions. Stmn2 levels in the media and islet extracts were measured using stathmin-2 ELISA kit (Cat# MBS7223765, MyBioSource) according to the supplier’s instruction. For each measurement, the values were compared between groups by t-test and among groups by one-way ANOVA (α = 0.05). Total cellular protein was determined using BCA assay and used to normalize cellular glucagon or Stmn2 per mg of cell protein. Then, alterations in values were expressed as percent changes compared to the baseline control. To this end, the Arg-stimulated secretion of glucagon or Stmn2 was normalized to the baseline level of glucagon and Stmn2, respectively, and expressed as relative fold changes (percent).

### Generation of diabetes mimicking αTC1-6 cells

Wild type αTC1-6 cells (a kind gift from C. Bruce Verchere, University of British Columbia, Vancouver, BC) were cultured in DMEM containing 16.7 mM glucose, L-glutamine, 15% horse serum and 2.5% fetal bovine serum, as described previously (Asadi and Dhanvantari, 2019; Asadi and Dhanvantari, 2020). As we have shown previously, after long-term culturing in high glucose conditions, cells demonstrated the phenotype of glucagon hypersecretion of diabetes(Asadi and Dhanvantari, 2019; McGirr et al., 2005).

### Inhibition of lysosomes in diabetes mimicking αTC1-6 cells and glucagon secretion studies

αTC1-6 cells were incubated for 24h in serum-free medium containing 16.7 mM glucose and 0.1% BSA. Cells were then washed twice with HBSS containing 16.7 mM glucose, HEPES, and 0.5% BSA and pre-incubated for 30 min in this medium. For lysosomal inhibition, cells were treated with 10 nM Bafilomycin A1 (Cat# B1793, Sigma) for 2 h. Cells were then rinsed and incubated in HBSS containing 16.7 mM glucose and 0.5% BSA for 2h. Secretion studies were done by addition of 55 mM KCl for 20 min as we have done previously (Asadi and Dhanvantari, 2020). Media were then collected into microfuge tubes containing PMSF (45 mM) and Aprotinin (5μg/mL), centrifuged at 14000×g for 15 minutes at 4°C, and then stored at −80°C until analysis. Cells were washed in ice-cold HBSS, lysed in ice-cold lysis buffer (50 mM Tris pH 7.4, 150 mM NaCl, 1% Triton X-100 plus 45 mM PMSF, and 5 μg/mL Aprotinin), and centrifuged at 14000×g for 15 minutes at 4°C. The supernatant was collected and stored at −80°C until analysis. Cell protein level was measured by BCA, as mentioned above, and used for normalization of glucagon secretion.

### Confocal immunofluorescences microscopy studies on diabetes mimicking αTC1-6 cells

Diabetes mimicking αTC1-6 cells were generated as mentioned above, and cultured on coverslips as described previously (Asadi and Dhanvantari, 2020). Experiments were done in the presence or absence of lysosomal inhibitor, BFA1, as described above. Cells were then cultured in serum-free DMEM containing 16.7 mM glucose and 0.1% BSA for 96h. Two hours before fixation, cells were incubated in HBSS containing 16.7 mM glucose and 0.5% BSA for 2h to ensure cells were treated the same as before secretion experiments. Cells were then washed 3X in PBS, fixed in 2% paraformaldehyde for 30 min, washed 5X in PBS, and incubated with blocking buffer (2% BSA in PBS containing 0.05% Tween 20) for 1h. Cells were then incubated with appropriate primary antibodies prepared in blocking buffer against glucagon (mouse anti-glucagon antibody, Cat # ab10988, Abcam; 1:1000 or rabbit anti-glucagon antibody, Cat# ab92517, Abcam; 1:25), Stathmin-2 (goat anti-SCG10 antibody, Cat # ab115513, Abcam; 1:250), or Lamp1 (mouse anti-Lamp1 antibody, Cat # ab25630, Abcam; 1:50). Following an overnight incubation at 4°C, coverslips were washed with PBS, and incubated for 2h in the dark at RT with appropriate secondary antibodies (donkey anti-mouse IgG Alexa Fluor 488, Cat# A-21202, Molecular Probes; donkey anti goat IgG Alexa Fluor 555, Cat# A-21432, Molecular Probes; donkey anti-rabbit IgG Alexa Fluor 647, Cat# A-31573, Molecular Probes). Then, coverslips were washed with PBS, stained with DAPI as previously described, and mounted on glass slides using ProLong antifade mountant. Image acquisition and analysis was done as described above. Experiments were repeated four times using freshly thawed cells.

### Statistical analyses

Comparison of values among groups was done by one-way ANOVA and between groups by t-test using Sigma Stat 3.5 software at α=0.05. For colocalization analysis of images, Pearson correlation coefficient values were extracted using NIS-Elements software and then values were compared between groups by t-test (α=0.05).

## Results

### Induction of diabetes in C57BL/6 mice

Measuring blood glucose levels following STZ injection showed fasting hyperglycemia (>14 mmol/L), indicating onset of diabetes (Figure 1A). Plasma levels of glucagon were significantly (p<0.001) higher and plasma levels of insulin were significantly reduced (p<0.001) in STZ-treated mice compared to the control group (Figure 1B, C), and there was a significantly (p<0.001) higher glucagon: insulin ratio in islets of STZ-treated mice compared to the control (Figure 1D). These findings indicate development of a diabetic metabolic and hormonal profile following STZ administration.

**Figure 1:**
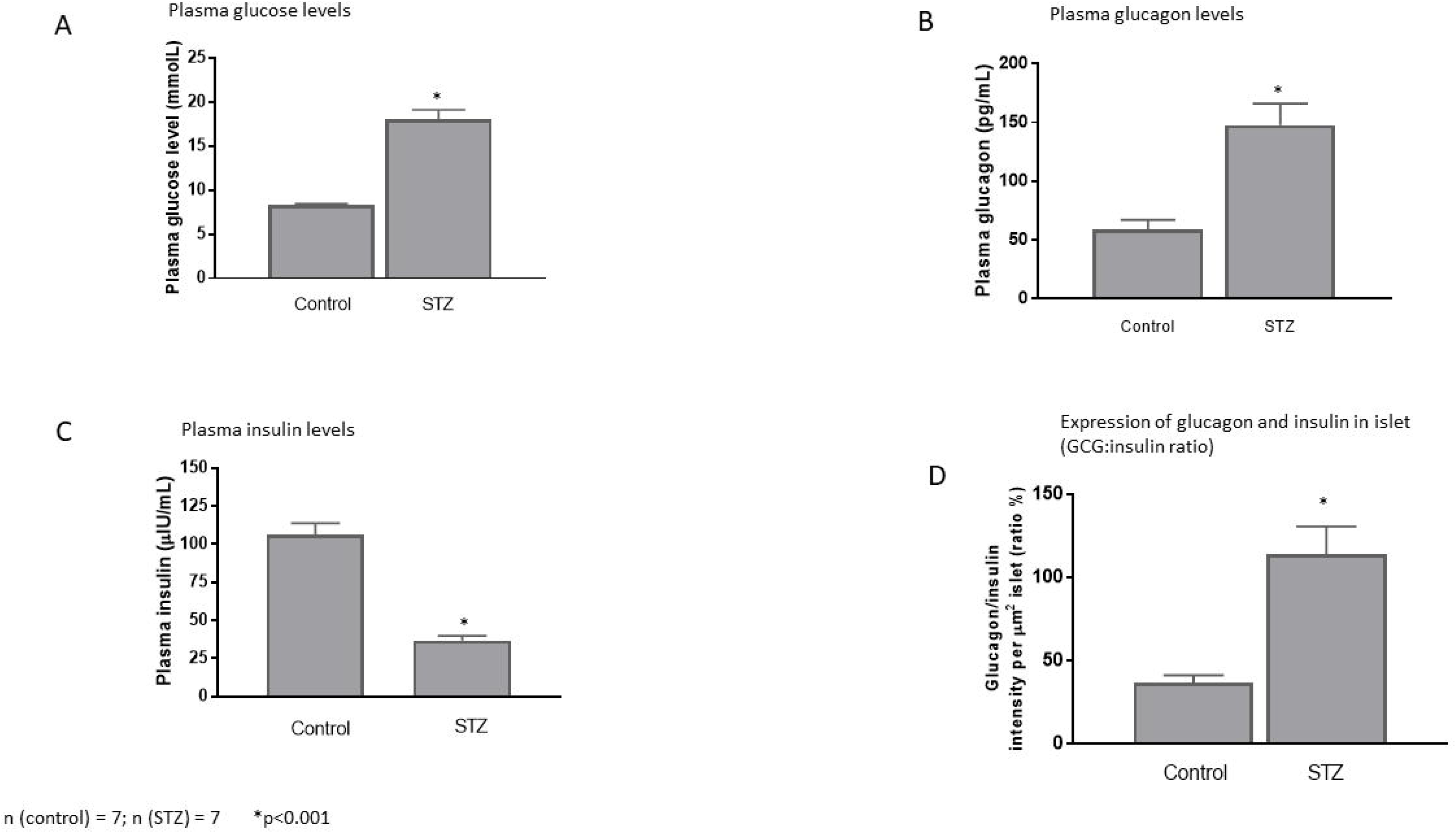
Streptozotocin (STZ) induced diabetes in C57BL/6 mice. (**A**) Plasma glucose, (**B**) plasma insulin, (**C**) plasma glucagon, and (**D**) glucagon: insulin ratio per μm^2^ of islet in STZ-induced diabetic mice. Values (mean± SD) were compared between diabetic (n=7) and control (n=7) mice by t-test. *p<0.001.

### Glucagon and Stmn2 colocalize in islets of diabetic and non-diabetic mice

Confocal immunofluorescence microscopy studies on islets of non-diabetic mice (Figure 2A) showed colocalization between glucagon and Stmn2 (Figure 2B), but not between insulin and Stmn2 (Figure 2C). This pattern of colocalization was conserved in islets from diabetic mice (Figures 2D-F). Quantification and analysis (Figure 2G) revealed strong colocalization between glucagon and Stmn2 in α-cells of both control (PCC 0.77± 0.02) and STZ-induced diabetic mice (PCC 0.83 ± 0.06). In contrast, there was almost no colocalization between insulin and Stmn2 in both control (PCC 0.04±0.03) and STZ-induced diabetic mice (PCC 0.03±0.06).

**Figure 2:**
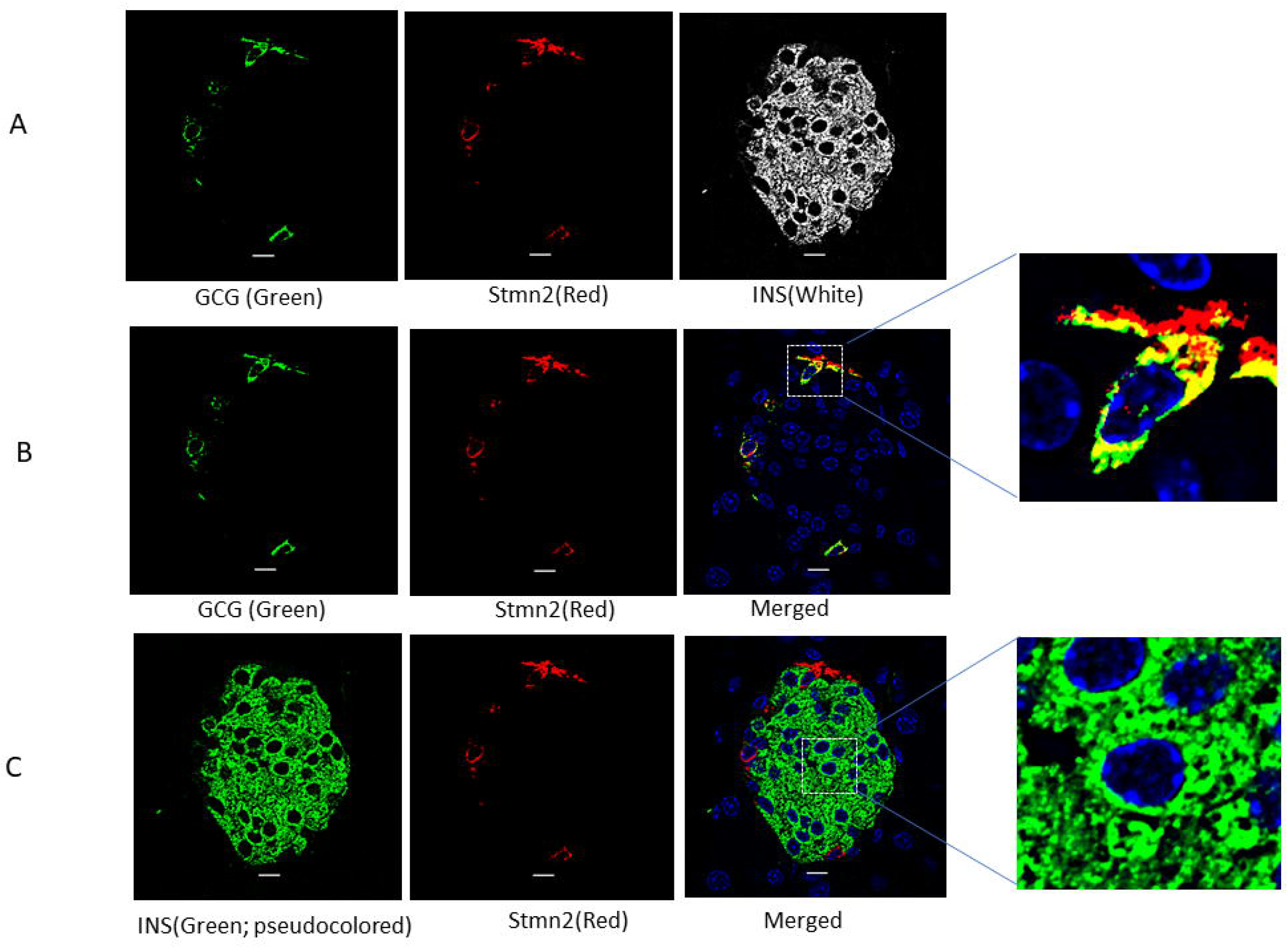
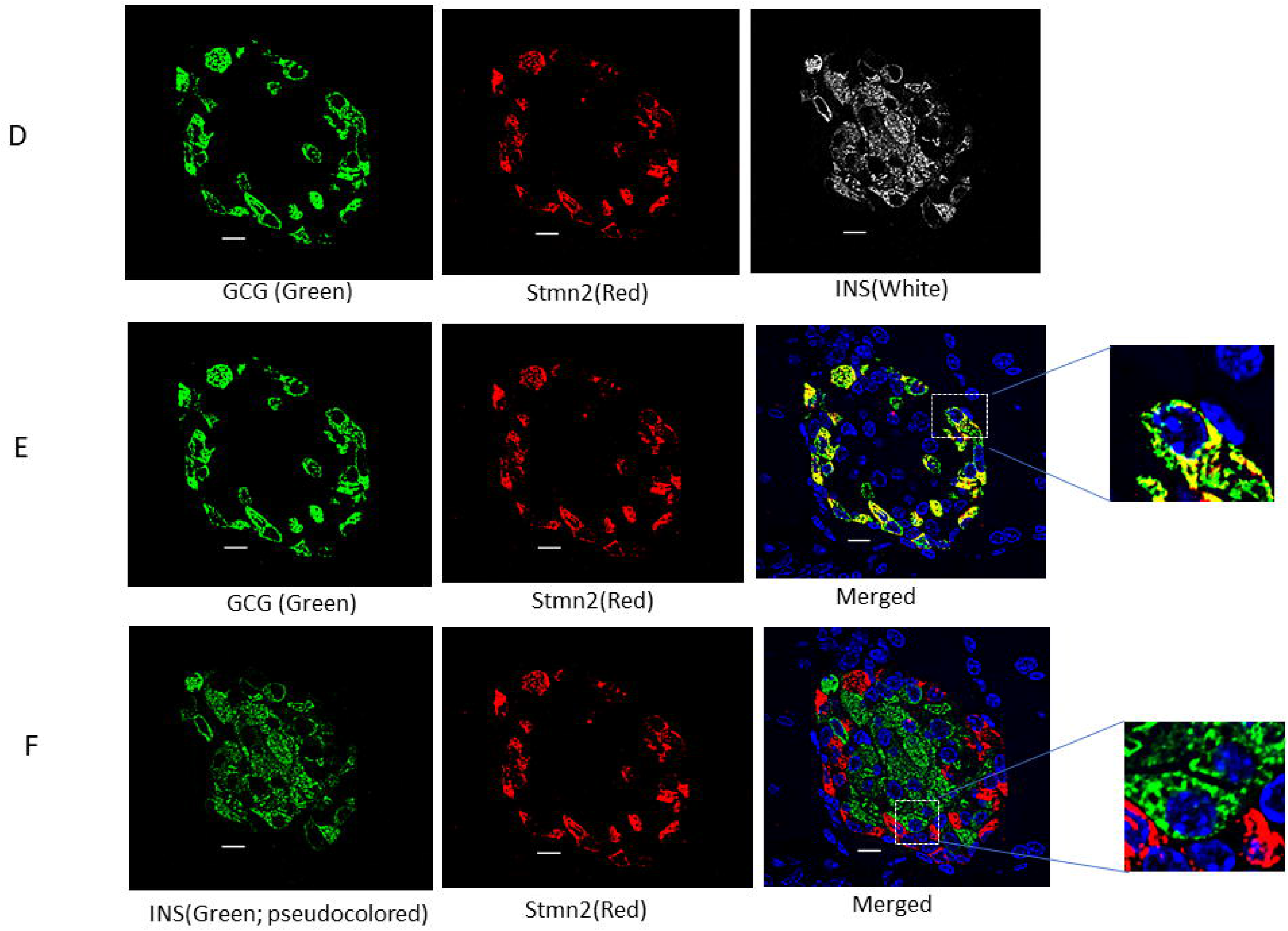
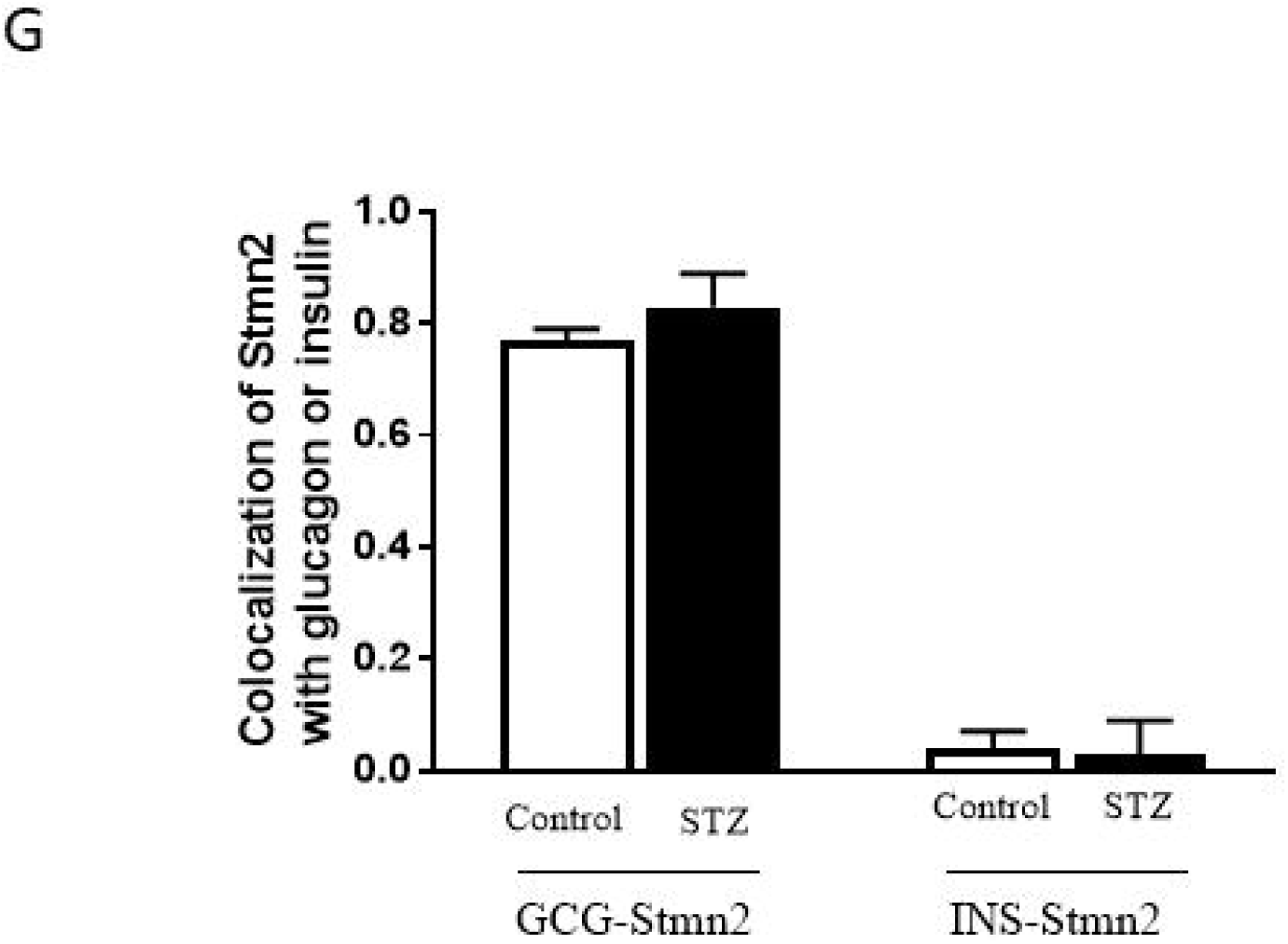
Stathmin-2 colocalizes with glucagon but not insulin in both non-diabetic and STZ-induced diabetic mice. (**A**) Islets of non-diabetic mice (n=7) were immunostained for glucagon (GCG), insulin (INS) and stathmin-2 (Stmn2); (**B**) after removing INS channel, colocalization was determined between GCG and Stmn2; (**C**) after removing GCG channel, and pseudocoloring of INS channel to green, colocalization was determined between Stmn2 and INS. **(D)** Islets of diabetic mice (n=7) were immunostained for GCG, INS and Stmn2; (**E**) after removing INS channel, colocalization was determined between GCG and Stmn2. (**F**) after removing GCG channel and pseudocoloring of INS channel to green, colocalization was determined between Stmn2 and INS. In each panel, marked area by square was magnified to show a typical individual cell. (**G**) Correlation between GCG and Stmn2 or INS and Stmn2 was determined by Pearson’s correlation coefficient (PCC) using NIS-Elements software.

### STZ-induced diabetes increases glucagon levels and reduces Stmn2 levels in α-cells

Analysis of fluorescence intensities revealed increased cellular levels of glucagon (p<0.001) and reduced levels of Stmn2 (p<0.01) in islets of STZ-induced diabetic mice (Figure 3A). As a consequence, the ratio of glucagon: Stmn2 in α-cells of STZ-induced diabetic mice significantly increased (p<0.01) compared to the non-diabetic controls (Figure 3B). Linear regression analysis of binary sum intensities showed a strong correlation between expression of Stmn2 and glucagon in α-cells of non-diabetic mice (R^2^ = 0.9, p<0.001) that was disrupted in STZ-induced diabetic mice (R^2^ = 0.07, p >0.05) (Figure 3C). This increase in cellular glucagon was paralleled by a ~ 4.5 times increase in the levels of *Gcg* mRNA, while there was no effect on *Stmn2* mRNA levels (Figure 4).

**Figure 3:**
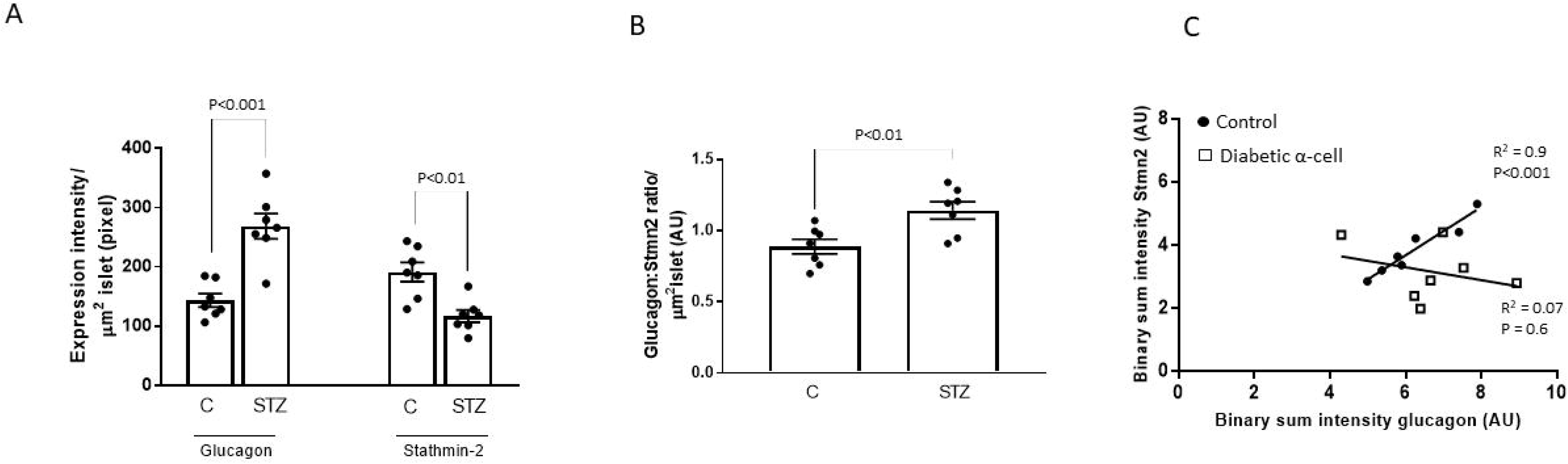
Altered ratios of stathmin-2 and glucagon in islets of STZ-induced diabetic mice. Islets were immunostained for stathmin-2 (Stmn2) and glucagon and images were acquired as described in Methods. (**A**) Expression of glucagon and Stmn2 were determined in islets of nondiabetic and diabetic mice by immunofluorescence intensity analysis. (**B**) Ratios of glucagon: Stmn2 levels were calculated per μm^2^ of islets in non-diabetic and diabetic mice. (**C**) Linear regression analysis on binary image intensities of the Stmn2 and glucagon. Filled circles and open squares demonstrate values in non-diabetic and diabetic islets, respectively.

**Figure 4:**
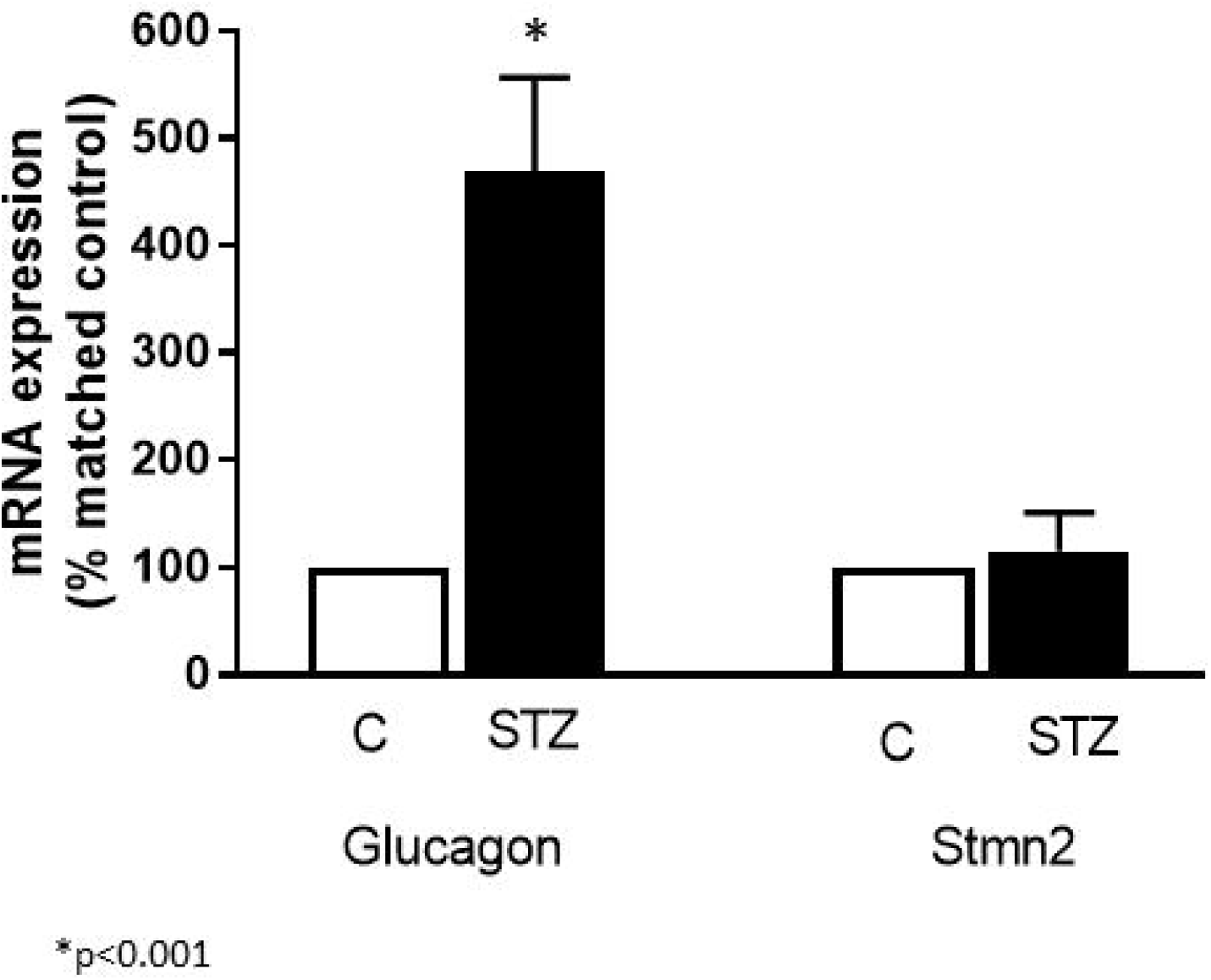
Expression of *Stmn2* and *Gcg* mRNA levels were determined in islets of non-diabetic (n=4) and diabetic (n=4) mice by qRT-PCR. Gene expression levels were normalized to that of *18S rRNA*. For each gene, alterations in the diabetic condition were normalized by the corresponding control group and expressed as percent of matched control. Comparison between control and STZ groups was done by t-test, α=0.05.

### Both glucagon and Stmn2 are localized within secretory granules of α-cells

Double immunogold-labeling transmission electron microscopy revealed colocalization of glucagon and Stmn2 within the secretory granule of mouse islet α-cells. Glucagon (10 nm particles; white arrows) and Stmn2 (18 nm particles; black arrows) were colocalized within the dense core of secretory granules in both non-diabetic (Figure 5A) and diabetic (Figure 5B) mouse α-cell.

**Figure 5:**
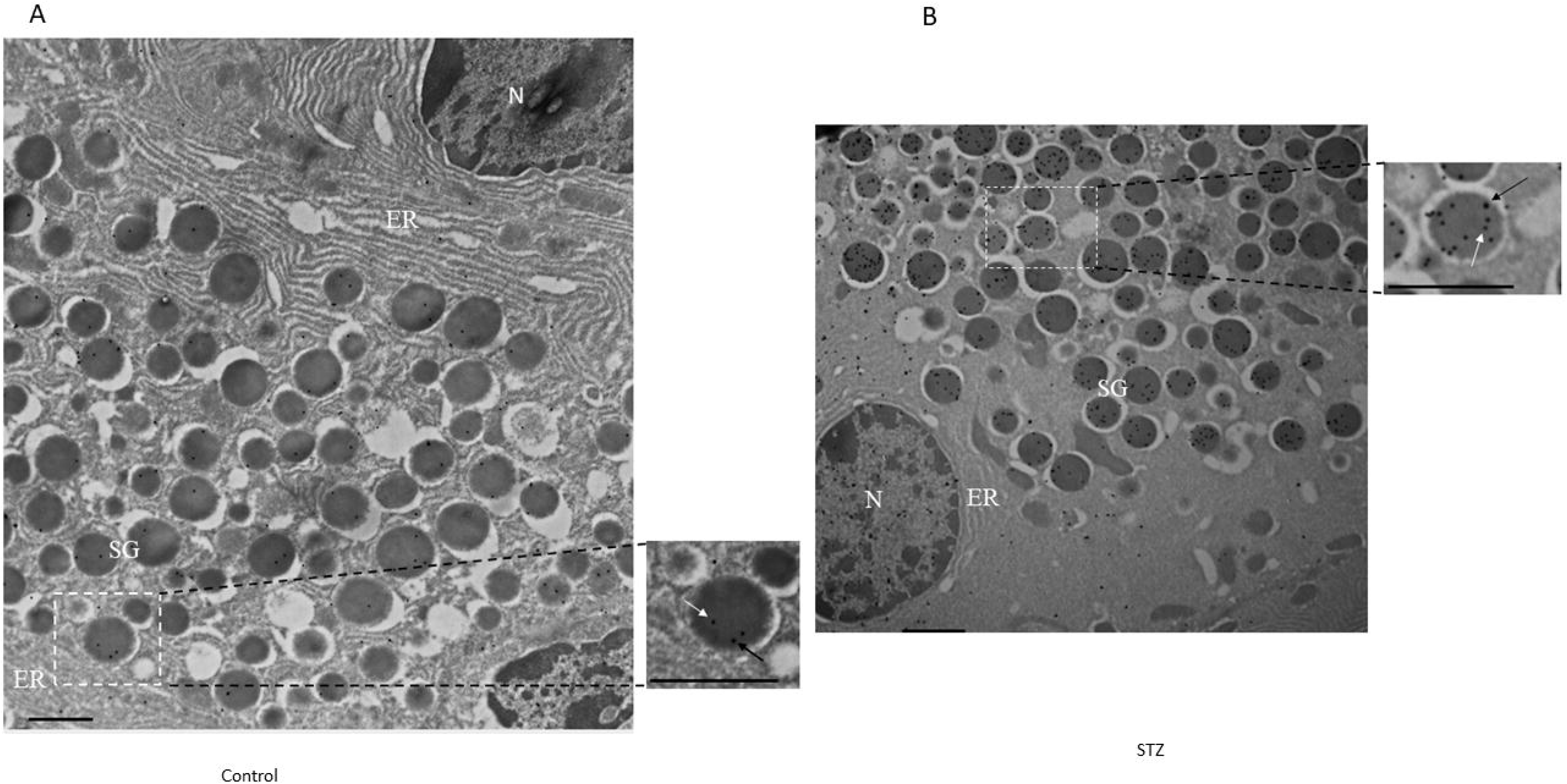
The presence of stathmin-2 and glucagon within secretory granules of α-cells. **(A)** Double immunogold labelling transmission electron microscopy of islets in non-diabetic and (**B**) STZ-induced diabetic mice was for glucagon (10 nm gold, white arrows in the magnified section) and stathmin-2 (18 nm gold, black arrows in the magnified section). The low magnification images (25000×, scale bar = 1 μm) show the ultrastructure of one α-cell. N (nucleus); ER (endoplasmic reticulum); SG (secretory granule); PM (plasma membrane). The magnified images (41000×, scale bar = 0.6 μm) highlight the presence of immunogold labels within a single secretory granule.

### Arginine stimulates parallel increases in glucagon and Stmn2 secretion

In line with our recent findings (Asadi and Dhanvantari, 2020), there was an increase in the secretion of both glucagon (~2.9 times) and Stmn2 (~2.3 times) from isolated islets in response to 25 mM Arg. In islets from diabetic mice, the glucagon secretory response was exaggerated, with increased basal and Arg-stimulated secretion (Figure 6A). There was also a small but significant increase in basal Stmn2 secretion and a significant increase in response to Arg in diabetic islets (Figure 6B). There was a concomitant reduction in cell glucagon content of non-diabetic islets in response to Arg (Figure 6A). In contrast, cell glucagon content of diabetic islets was elevated in the absence of Arg, and remained elevated after Arg stimulation (Figure 6A), consistent” with the diabetic phenotype of glucagon production. In parallel with the pattern of cell glucagon content in nondiabetic islets, cell Stmn2 content also decreased in response to Arg (Figure 6B). Importantly, cell Stmn2 content was reduced in islets of diabetic mice, and further reduced after Arg stimulation (Figure 6B), thereby showing a different profile from that of glucagon in diabetes.

**Figure 6:**
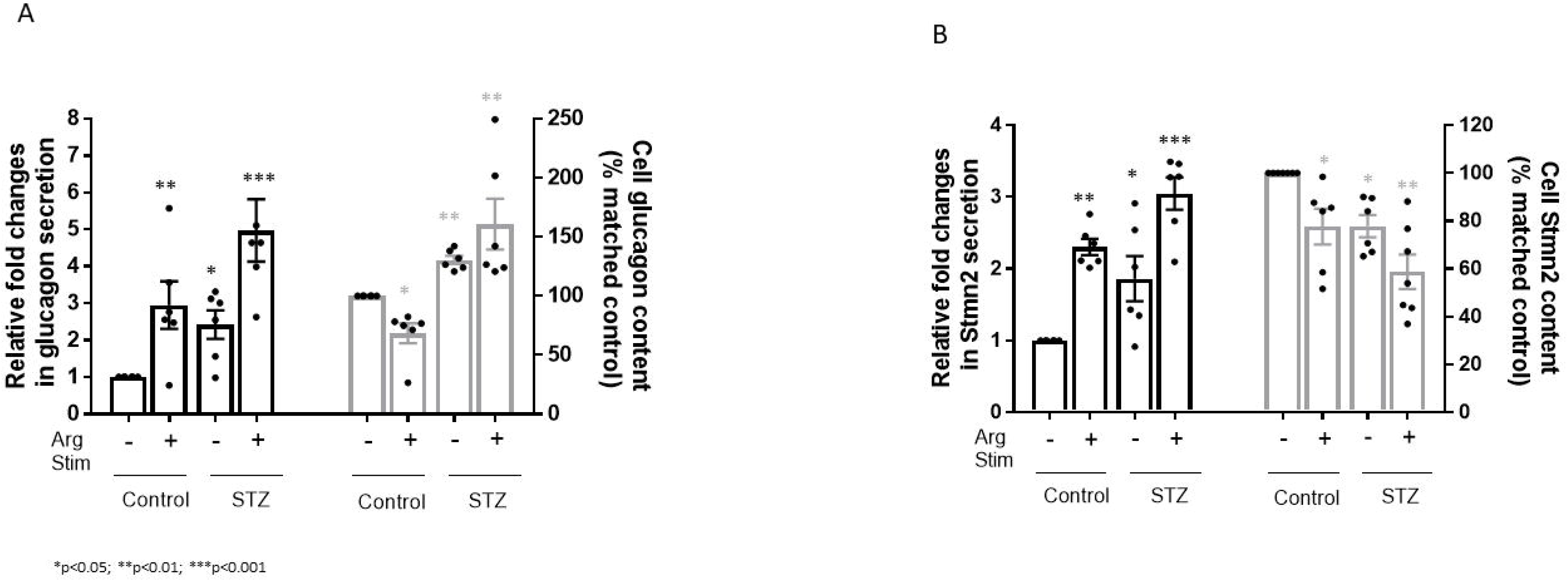
Parallel alterations in secretion of glucagon and Stmn2 from α-cells in both nondiabetic and diabetic mice. **(A)** Secretion of glucagon and islet glucagon content in isolated islets of non-diabetic (C; n=7) and diabetic mice (STZ; n=7) at the presence or absence of Arginine (25 mM, 20 min). Secretion values were normalized by baseline control secretion and expressed as fold changes. Glucagon contents were normalized by baseline control and expressed as percent changes. **(B)** Secretion of Stmn2 and islet Stmn2 content in isolated islets of non-diabetic (C; n=7) and diabetic (STZ; n=7) mice in the presence or absence of Arginine (25 mM, 20 min). Secretion values were normalized by baseline secretion in control and expressed as fold changes. Glucagon content was normalized by baseline content in control and expressed as percent changes. Values were expressed as mean ± SEM and compared among groups by 1-Way ANOVA at α=0.05. *p<0.05; **p<0.01; ***p<0.001.

### Trafficking of glucagon and Stmn2 towards the Lamp2A^+^ lysosome is inhibited in islets from STZ-induced diabetic mice

Based on our recent study that indicated a role for Stmn2 in regulating glucagon secretion by trafficking through the endolysosomal compartment in αTC1-6 cells (Asadi and Dhanvantari, 2020), we were interested to determine if the diabetes-induced alterations in the levels of Stmn2 and glucagon in mouse islets were associated with changes in the pattern of intracellular trafficking through the endolysosomal system. We therefore determined the presence of glucagon and Stmn2 in all four compartments of the endolysosomal pathway (the lysosome, late endosome, early endosome, and recycling endosome) in normal and diabetic mouse islets.

Using confocal immunofluorescence microscopy (Figure 7A), we identified individual α-cells in which Lamp2A colocalized with either glucagon (Figure 7B) or Stmn2 (Figure 7C) in normal mouse islets. In contrast, Lamp2A did not colocalize with either glucagon or Stmn2 in individual α-cells in islets of STZ-induced diabetic mice (Figures 7D-F). Quantification and analysis of colocalization showed a moderate level of colocalization between Lamp2A and glucagon (PCC 0.48 ± 0.08) and between Lamp2A and Stmn2 (PCC 0.52 ± 0.09) in the control group (Figure 7G). In contrast, STZ-induced diabetes significantly reduced levels of colocalization between glucagon and Lamp2A (PCC 0.14 ± 0.03; p<0.001) and also between Stmn2 and Lamp2A (PCC 0.15 ± 0.03; p<0.001) (Figure 7G).

**Figure 7:**
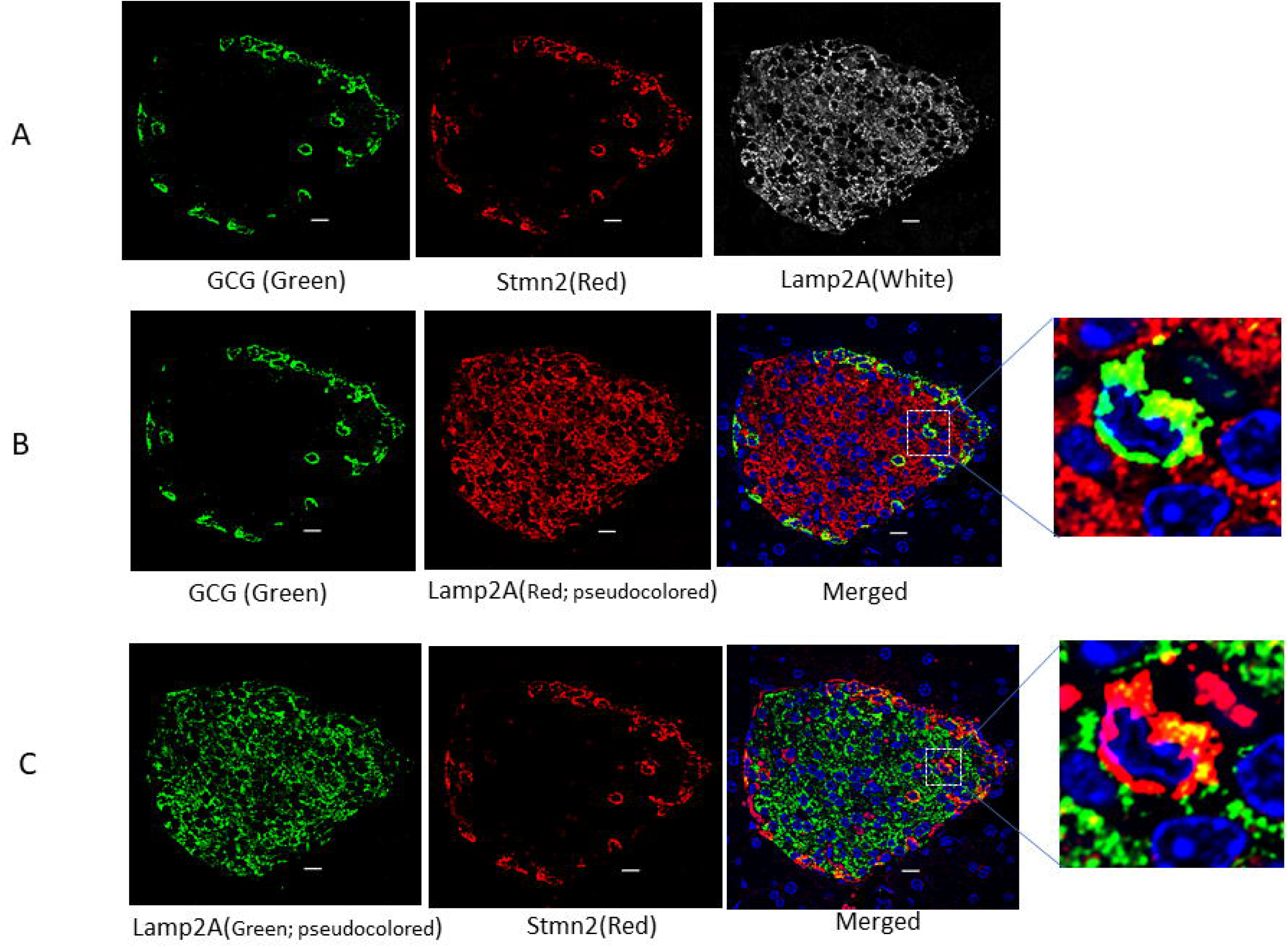
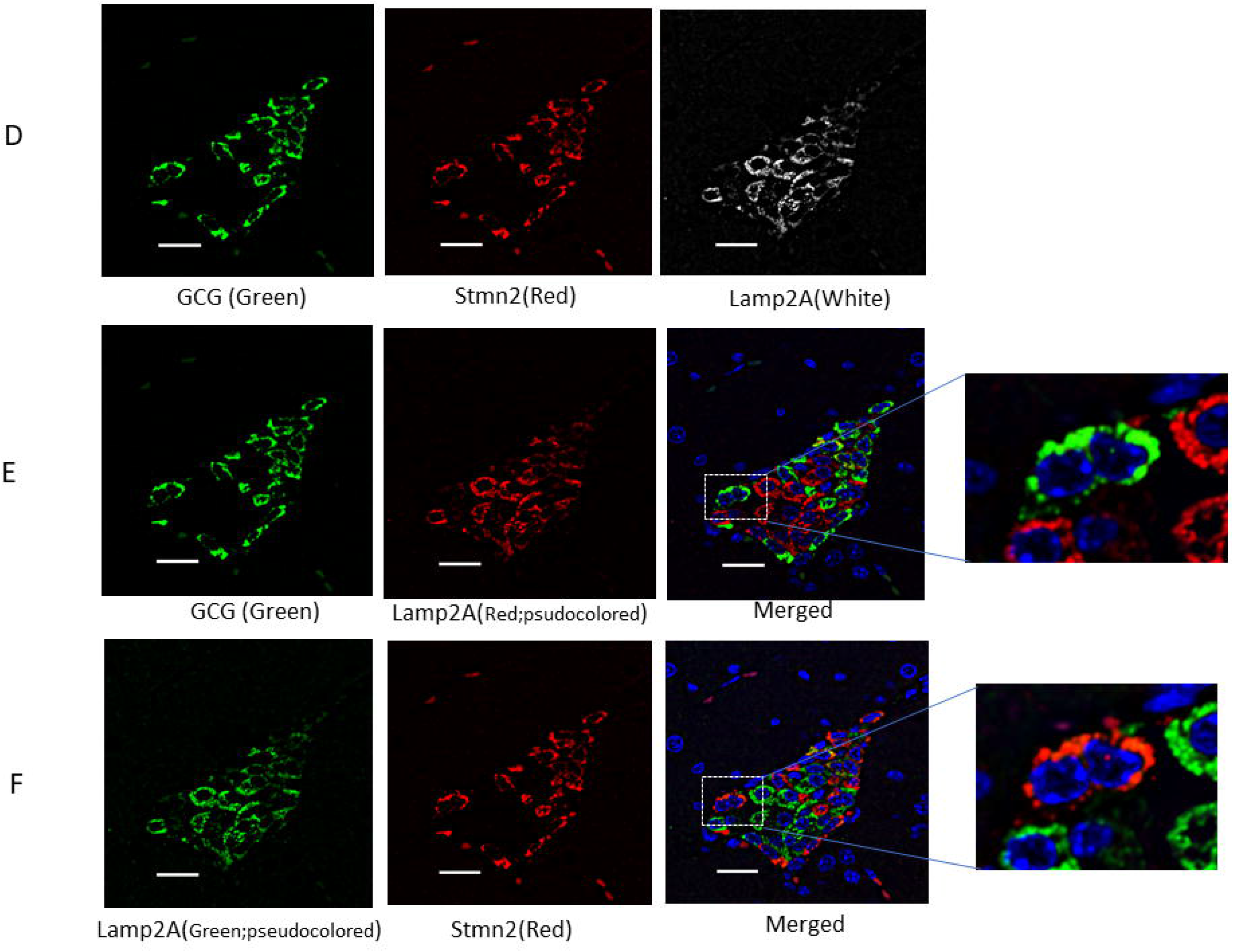
The localization of Stmn2 and glucagon in Lamp2A^+^ lysosomes of α-cells is inhibited in diabetes. **(A)** Islets from non-diabetic mice (n=7) were immunostained with antibodies against glucagon (GCG), stathmin-2 (Stmn2) and the lysosomal marker, Lamp2A. Representative images are shown. **(B)** Colocalization of glucagon and Lamp2A. The Lamp2A image was pseudocoloured red for visualization of colocalization in the merged image and inset. **(C)** Colocalization of Stmn2 and Lamp2A. The Lamp2A image was pseudocoloured green for visualization of colocalization in the merged image and inset. **(D)** Islets of diabetic mice (n=7) were immunostained for glucagon, Stmn2 and Lamp2A. **(E)** Colocalization of glucagon and Lamp2A. The Lamp2A image was pseudocoloured red for visualization of colocalization in the merged image and inset. **(F)** Colocalization of Stmn2 and Lamp2A. The Lamp2A image was pseudocoloured green for visualization of colocalization in the merged image and inset. All images were acquired and post-processed as described in Methods. In each merged panel, selected areas (white square) were magnified to show individual cells within islets. **(G)** Analysis of colocalization of glucagon and LAMP2A, and Stmm2 and LAMP2A in normal and diabetic (STZ) islets. Pearson’s correlation coefficient (PCC) values are shown as means ± SEM. Each dot represents a mean of 15 images per mouse.

### STZ-induced diabetes increased the localization of Stmn2 in late endosomes

The late endosome marker, Rab7, colocalized with glucagon (Figure 8B), but did not appear to colocalize with Stmn2 (Figure 8C) in non-diabetic α-cells. However, following induction of diabetes, Rab7 did appear to colocalize with Stmn2 (Figure 8F) and maintained colocalization with glucagon (Figure 8E). Quantification and analysis showed a moderate level of colocalization between glucagon and Rab7 in islets of both control (PCC 0.42 ± 0.1) and STZ-induced diabetic (PCC 0.48±0.1) mice (Figure 8G). Colocalization between Stmn2 and Rab7 in the control group was weak (PCC 0.29 ± 0.12), but significantly increased (PCC 0.48 ± 0.09, p<0.01) in islets from STZ-induced diabetic mice.

**Figure 8:**
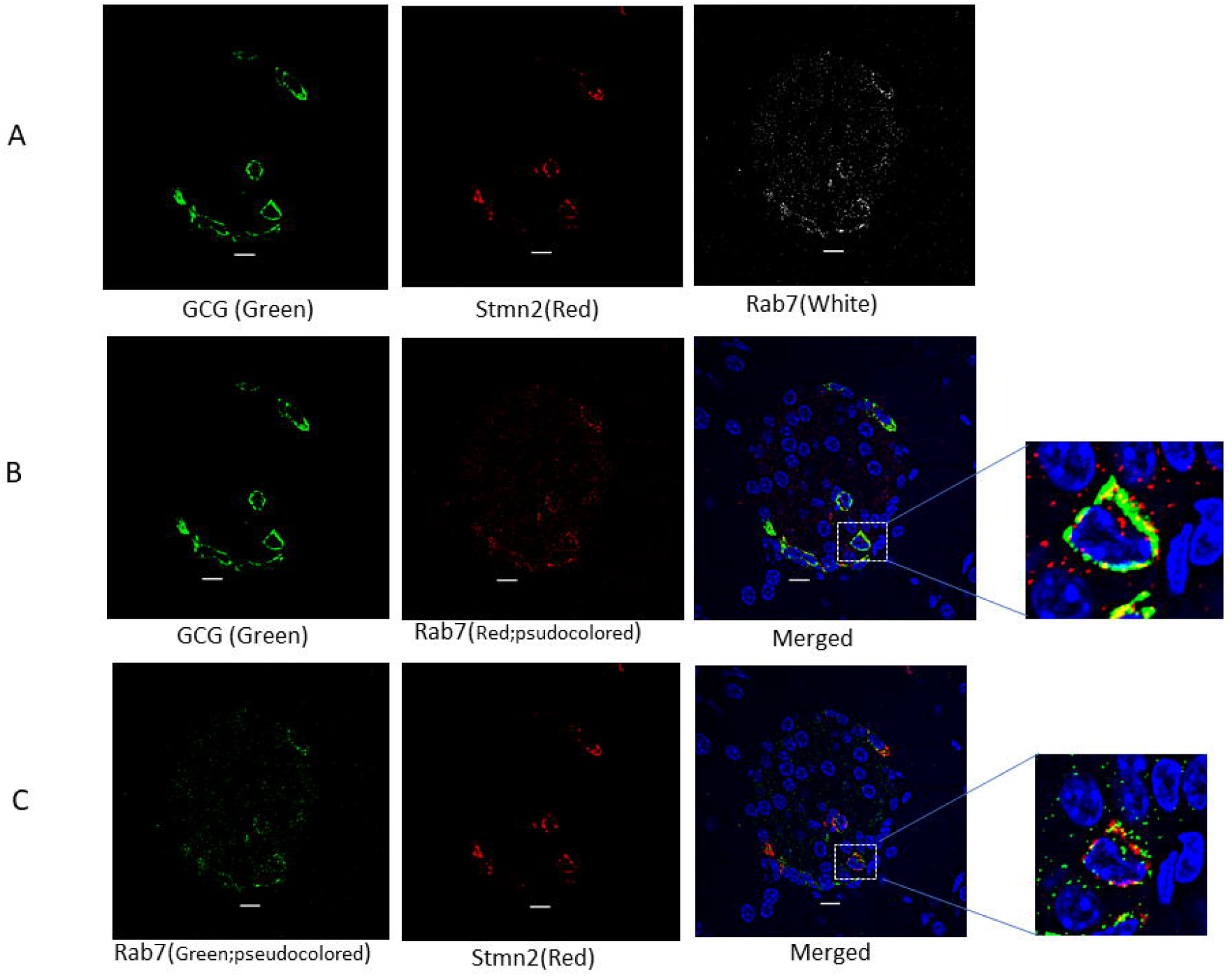
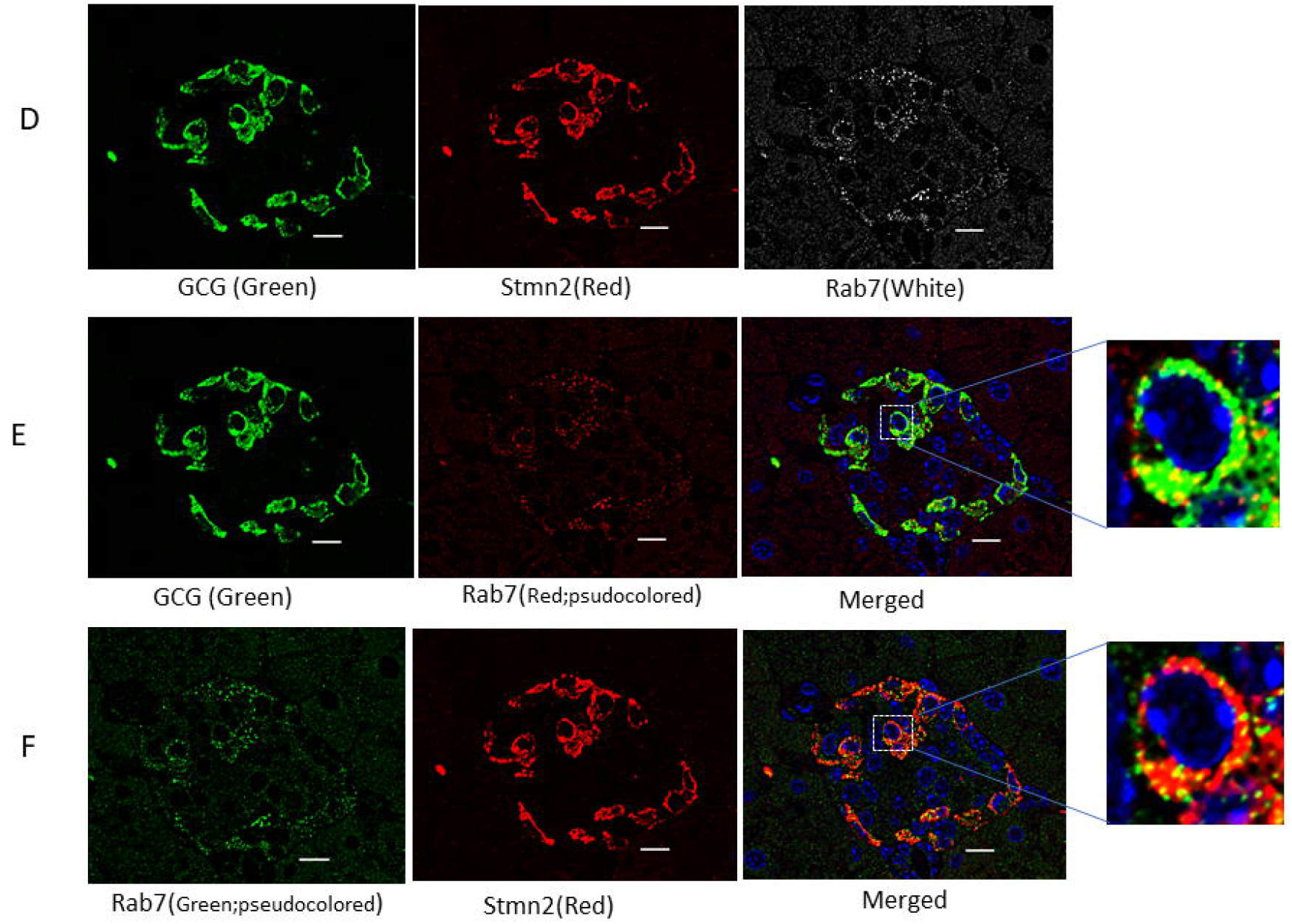
Diabetes enhances colocalization of Stmn2 with Rab7^+^ late endosomes in α-cells. **(A)** Islets from non-diabetic mice (n=7) were immunostained with antibodies against glucagon (GCG), stathmin-2 (Stmn2) and the late endosome marker, Rab7. Representative images are shown. **(B)** Colocalization of glucagon and Rab7. The Rab7 image was pseudocoloured red for visualization of colocalization in the merged image and inset. **(C)** Colocalization of Stmn2 and Rab7. The Rab7 image was pseudocoloured green for visualization of colocalization in the merged image and inset. **(D)** Islets from diabetic mice (n=7) were immunostained for glucagon, Stmn2 and Rab7; **(E)** Colocalization of glucagon and Rab7. The Rab7 image was pseudocoloured red for visualization of colocalization in the merged image and inset. **(F)** Colocalization of Stmn2 and Rab7. The Rab7 image was pseudocoloured green for visualization of colocalization in the merged image and inset. All images were acquired and post-processed as described in Methods. In each merged panel, selected areas (white square) were magnified to show individual cells within islets. **(G)** Analysis of colocalization of glucagon and Rab7, and Stmm2 and Rab7 in normal and diabetic (STZ) islets. Pearson’s correlation coefficient (PCC) values are shown as means ± SEM. Each dot represents a mean of 15 images per mouse. **(H)** Fluorescent intensities of glucagon, insulin (INS) and Rab7 were shown in islets from nondiabetic and **(I)** diabetic mice. **(J)** Analysis of fluorescent intensities of glucagon, insulin and Rab7 in α and β-cells. Intensities of DAPI-stained nuclei were used to normalize Rab7 intensities.

### Glucagon and Stmn2 do not localize within the early endosome

The early endosome marker, EEA1, appeared to be localized strongly to the core of the islet (Figure 9A). As shown by the magnified images, EEA1 did not colocalize with either glucagon (Figure 9B) or Stmn2 (Figure 9C). These patterns of colocalization remained unchanged in diabetes (Figures 9D-F). Quantification and analysis showed a very weak colocalization of EEA1 with glucagon (PCC 0.07 ± 0.05) or Stmn2 (PCC 0.07 ± 0.04) in the control group. Following induction of diabetes, colocalization of EEA1 with glucagon (PCC 0.03 ± 0.07) or Stmn2 (PCC 0.04 ± 0.07) still remained very weak.

**Figure 9:**
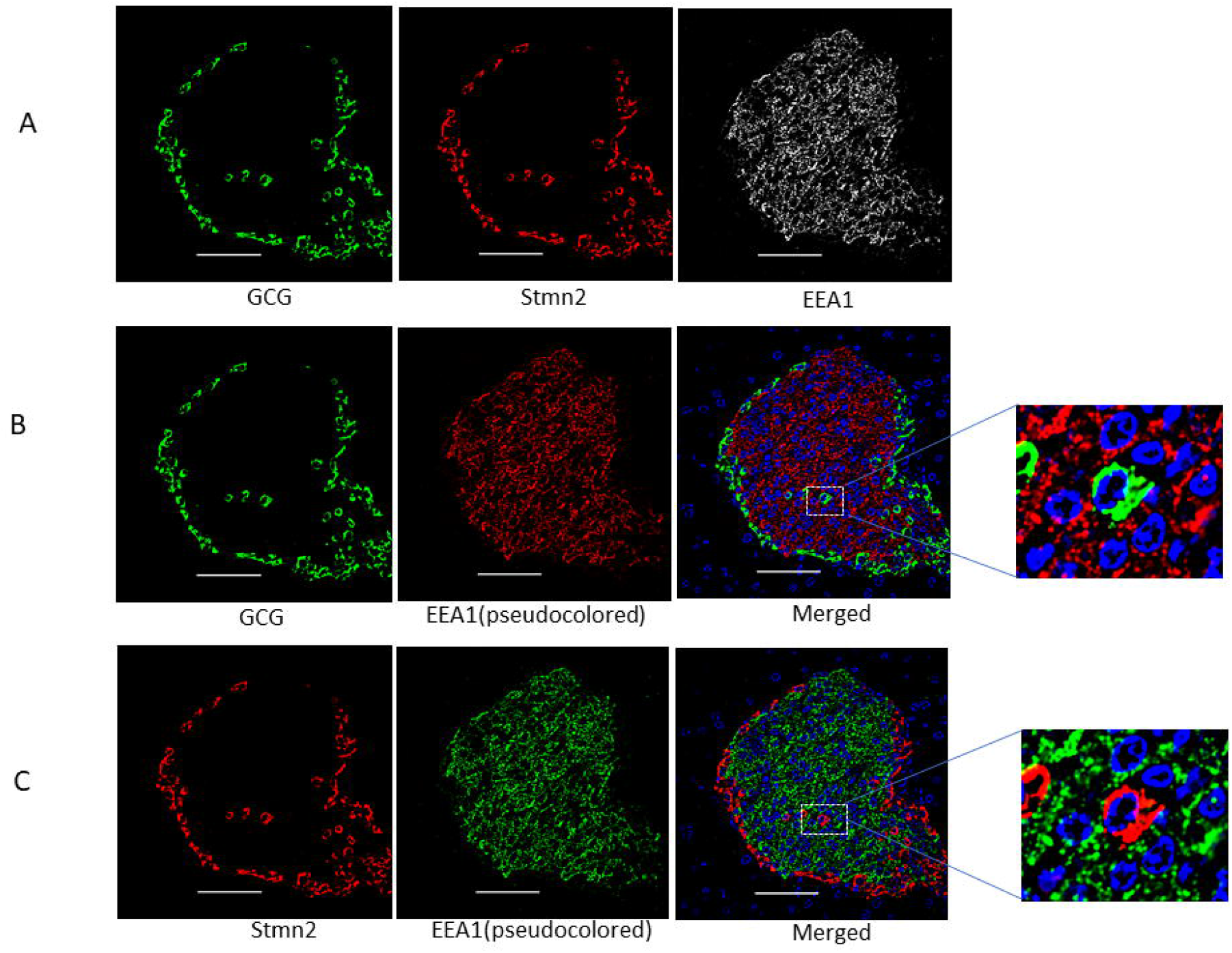
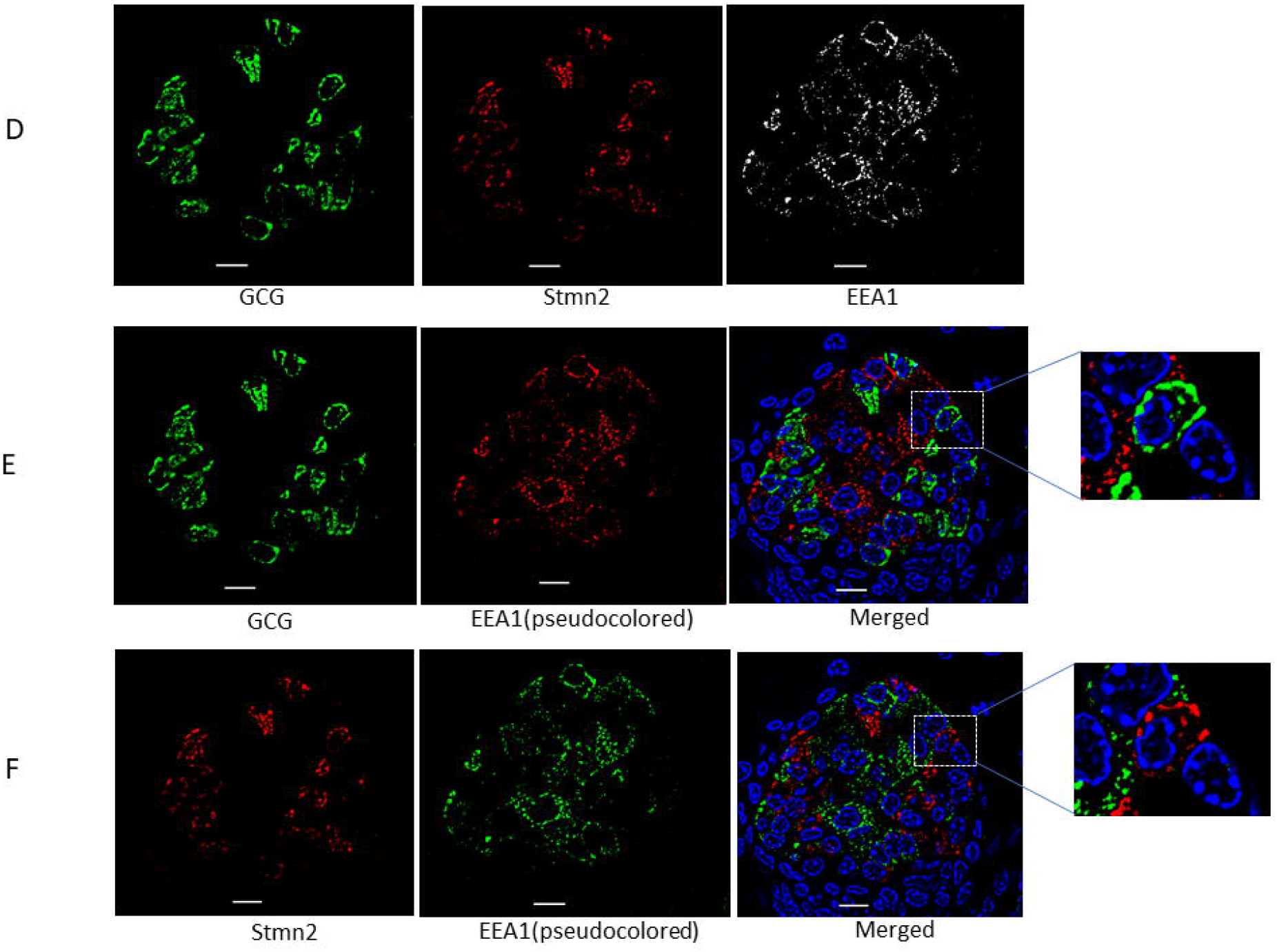
Glucagon and stathmin-2 are not localized in early endosomes in α-cells. **(A)** Islets from non-diabetic mice (n=7) were immunostained with antibodies against glucagon (GCG), stathmin-2 (Stmn2) and the early endosome marker, EEA1. Representative images are shown. **(B)** Colocalization of glucagon and EEA1. The EEA1 image was pseudocoloured red for visualization of colocalization in the merged image and inset. **(C)** Colocalization of Stmn2 and EEA1. The EEA1 image was pseudocoloured green for visualization of colocalization in the merged image and inset. **(D)** Islets from diabetic mice (n=7) were immunostained for glucagon, Stmn2 and EEA1. **(E)** Colocalization of glucagon and EEA1. The EEA1 image was pseudocoloured red for visualization of colocalization in the merged image and inset. **(F)** Colocalization of Stmn2 and EEA1. The EEA1 image was pseudocoloured green for visualization of colocalization in the merged image and inset. All images were acquired and post-processed as described in Methods. In each merged panel, selected areas (white squares) were magnified to show individual cells within islets. **(G)** Analysis of colocalization of glucagon and EEA1, and Stmm2 and EEA1 in normal and diabetic (STZ) islets. Pearson’s correlation coefficient (PCC) values are shown as means ± SEM. Each dot represents a mean of 15 images per mouse.

### Presence of glucagon in recycling endosome is not prominent

The immunofluorescence signal of the recycling endosome marker, Rab11A, also appeared to be quite strong in the core of the islet (Figure 10A), and did not colocalize with glucagon (Figure 10B), but colocalized with insulin (Figure 10C). As well, STZ-induced diabetes did not alter the pattern of distribution between Rab11A and glucagon (Figure 10D), but did decrease the colocalization between insulin and Rab11A (Figures 10D-F). In addition, another recycling endosome marker, Rab11B, showed a similar distribution within the islet (Figure 10 G); it also did not colocalize with glucagon (Figure 10H) but colocalized with insulin (Figure 10I); this pattern remained unchanged in diabetes (Figures 10K, L). Quantification and analysis (Figure 10M) confirmed the weak colocalization between glucagon and Rab11A (PCC 0.13 ± 0.05) or Rab11B (PCC 0.13 ± 0.04) in non-diabetic mice, and in diabetic mice (colocalization of glucagon with Rab11A, PCC 0.13 ± 0.04 or Rab11B, PCC 0.13 ± 0.03). In contrast, insulin showed a strong colocalization with Rab11A (PCC 0.63 ± 0.09) or Rab11B (PCC 0.61 ± 0.06), which significantly decreased (p<0.001) following induction of diabetes (colocalization of insulin with Rab11A, PCC 0.35 ± 0.5, or Rab11B, PCC 0.38 ± 0.07).

**Figure 10:**
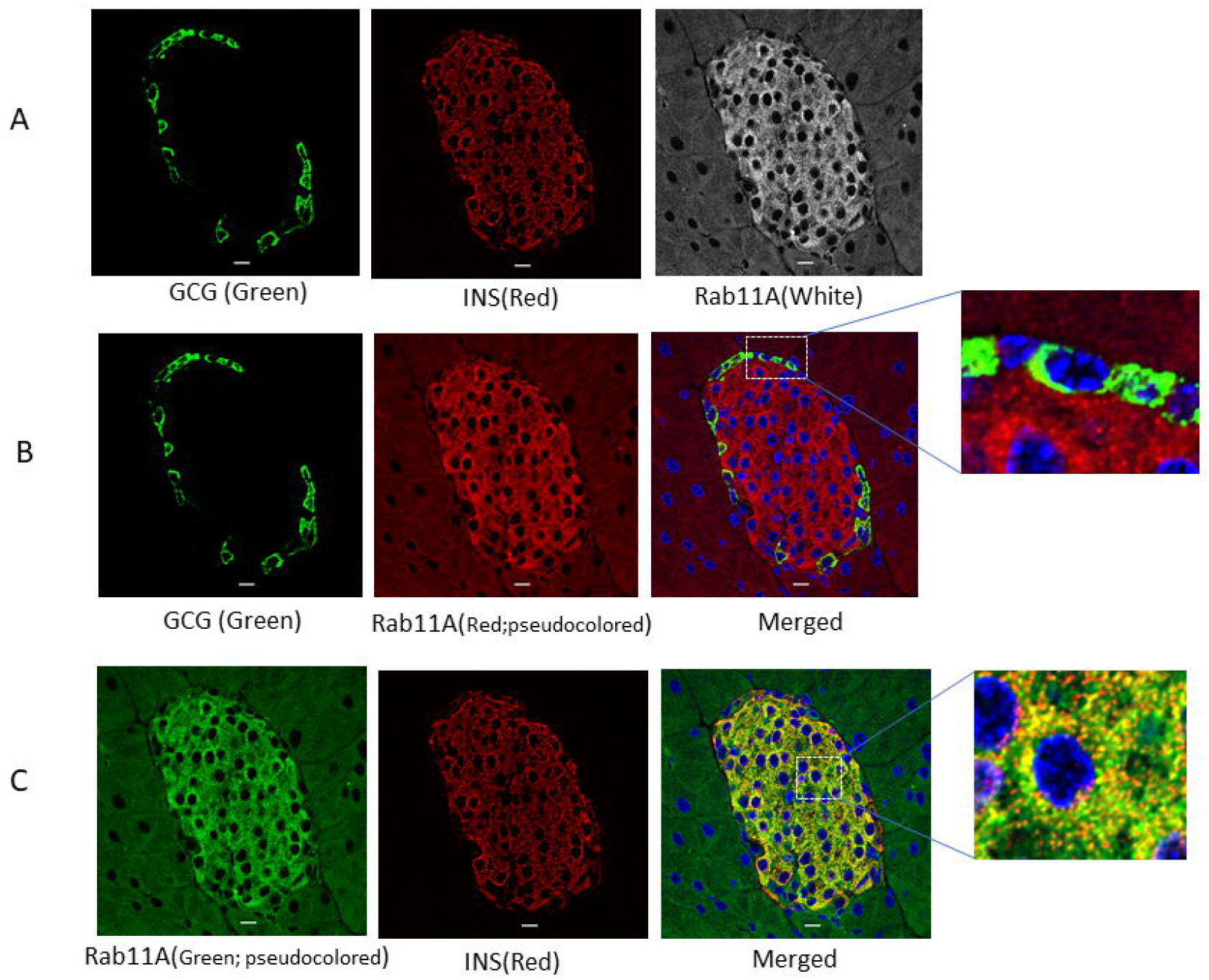
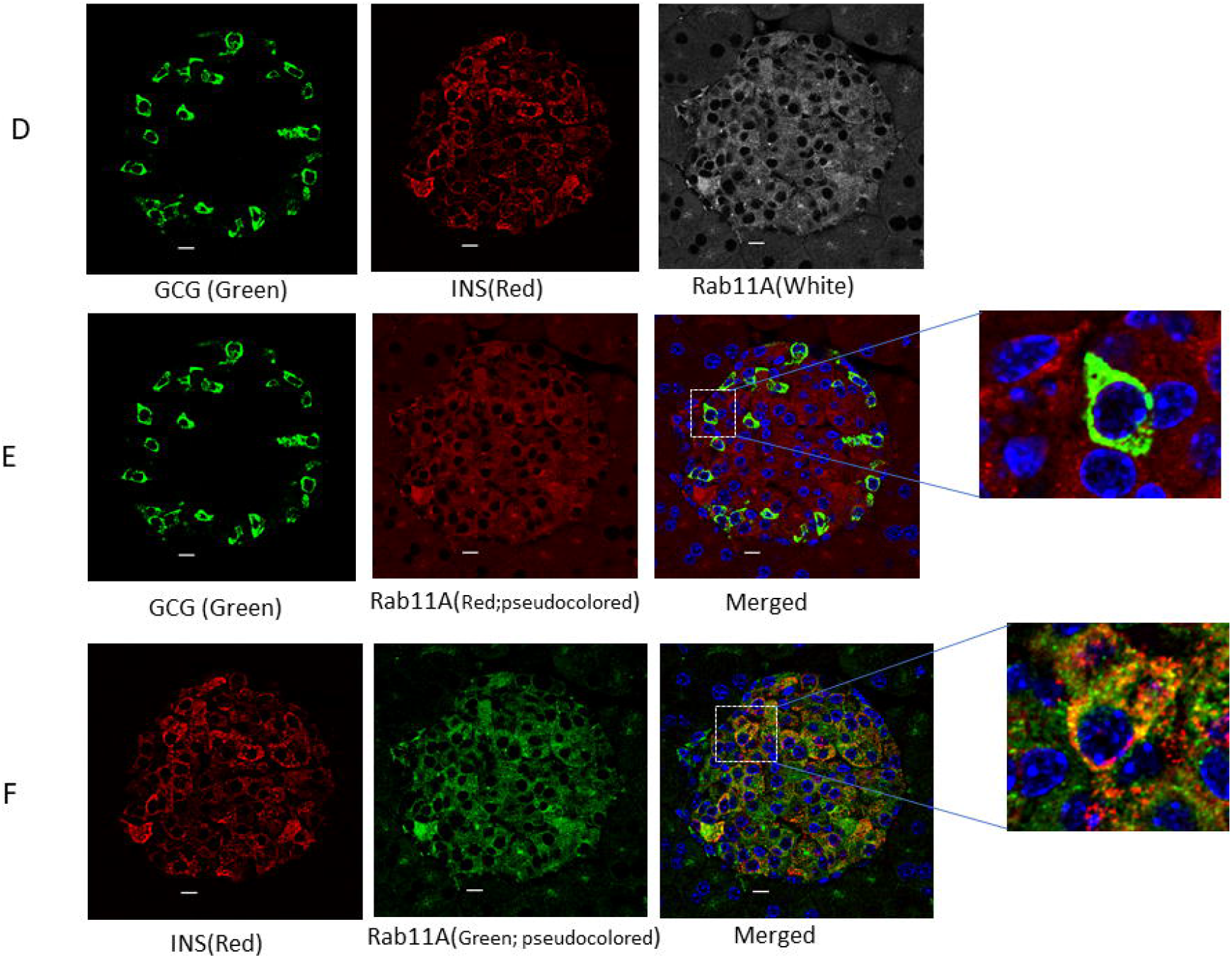
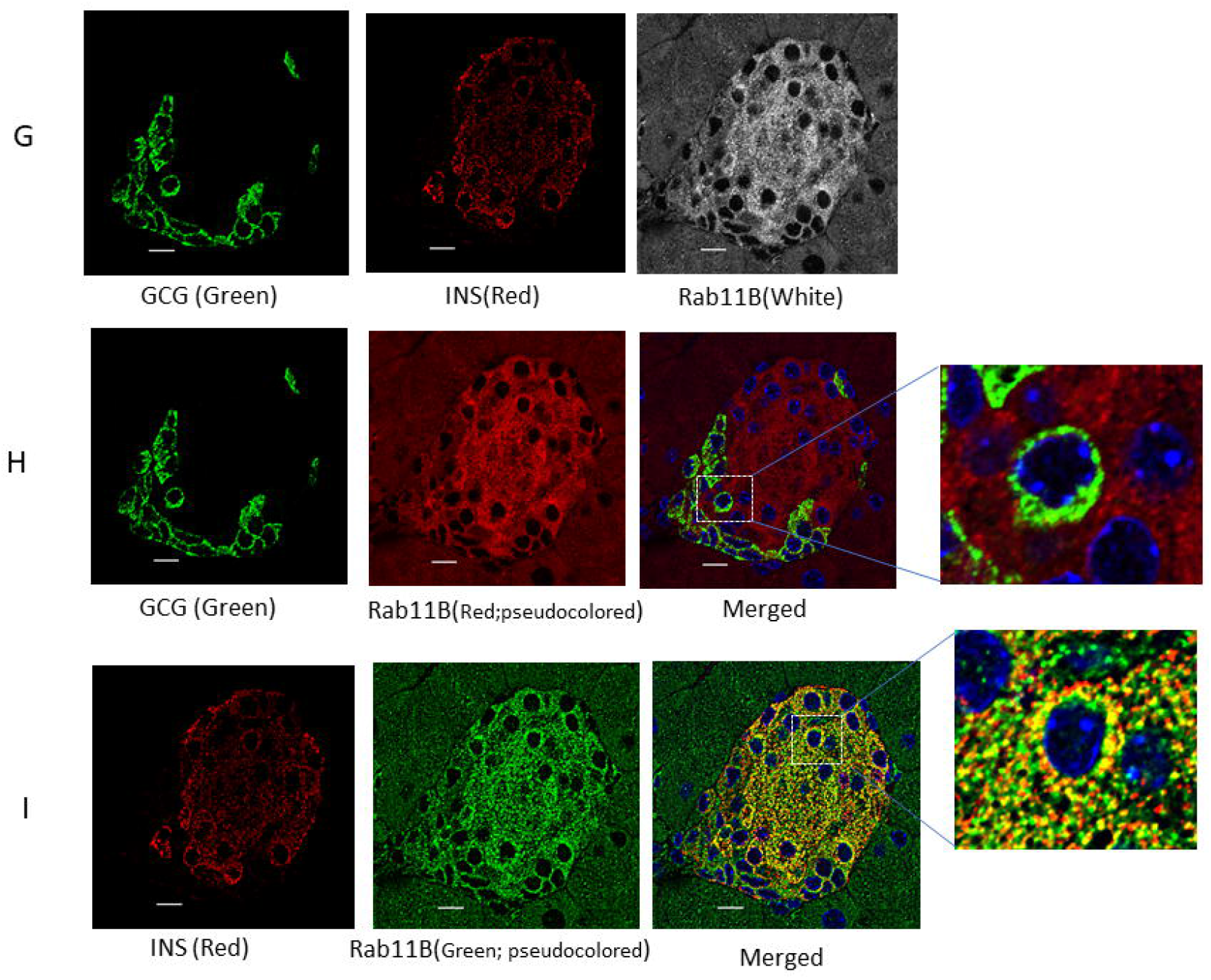
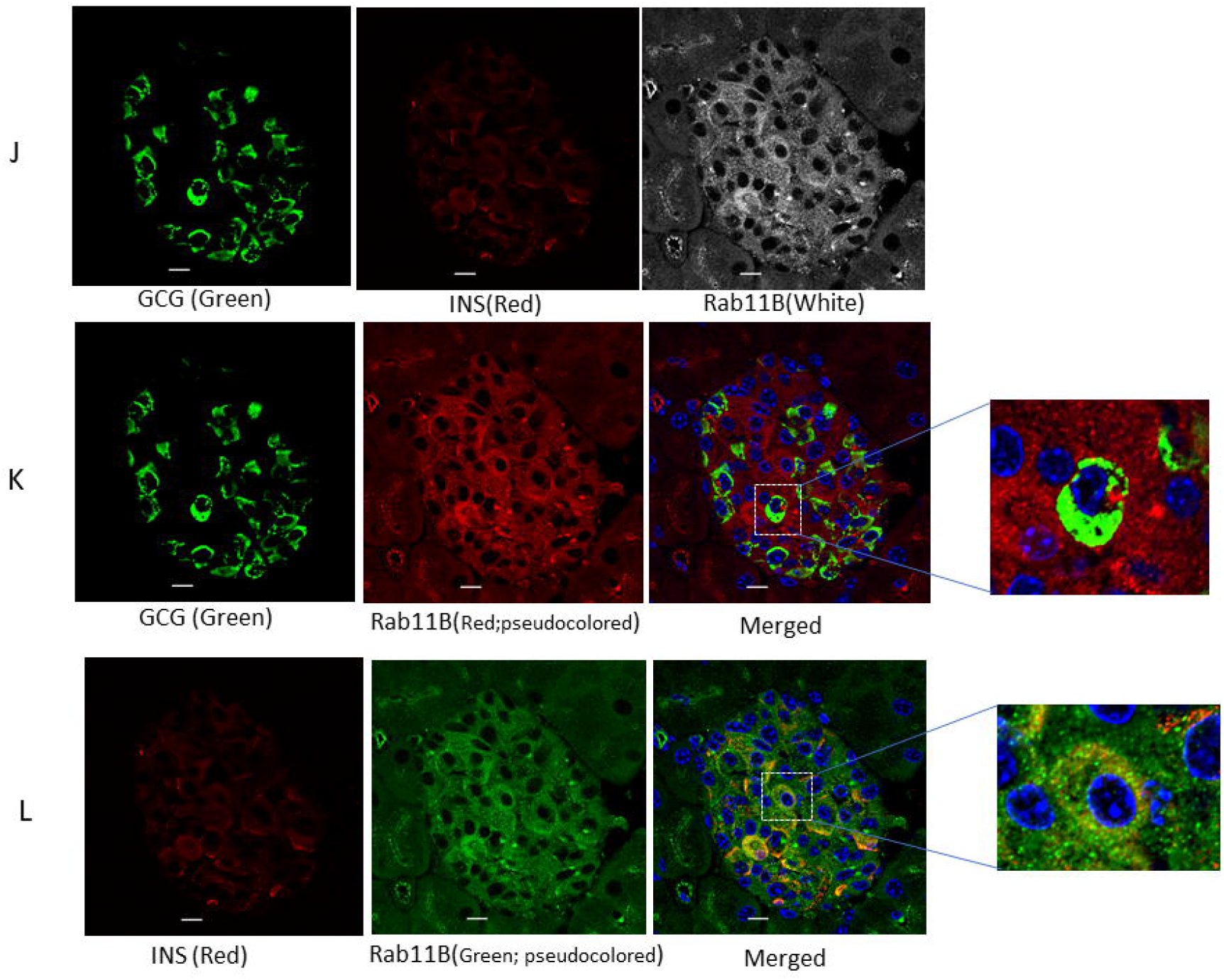
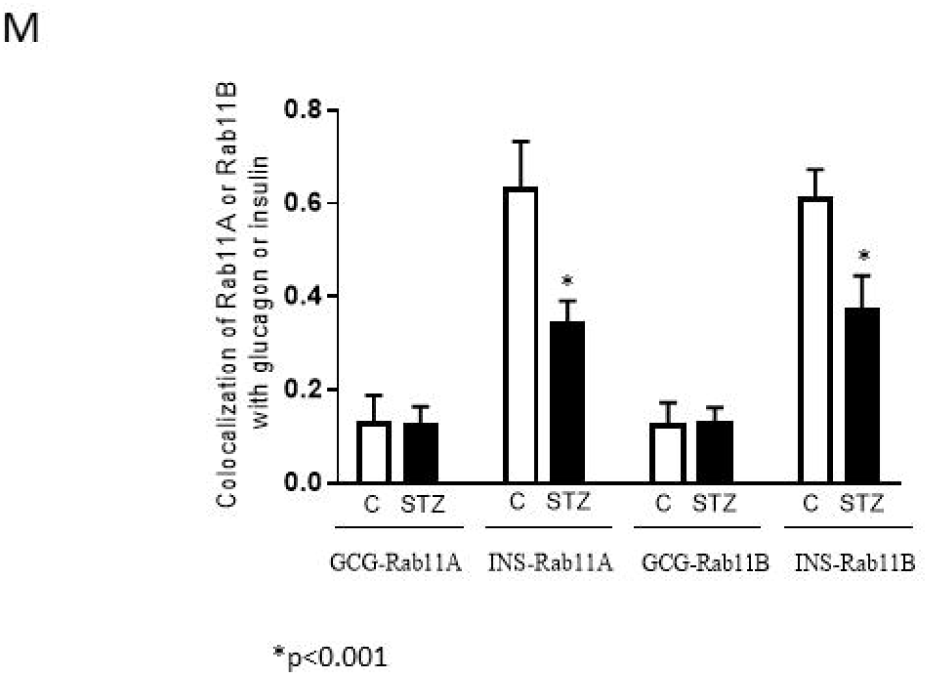
Glucagon is not localized in recycling endosome of α-cells. (**A**) Islets from nondiabetic mice (n=7) were immunostained with antibodies against glucagon (GCG), insulin (INS) and the recycling endosome marker, Rab11A. **(B)** Colocalization of glucagon and Rab11A. The Rab11A image was pseudocoloured red for visualization of colocalization in the merged image and inset. **(C)** Colocalization of insulin and Rab11A. The Rab11A image was pseudocoloured green for visualization of colocalization in the merged image and inset. **(D)** Islets from diabetic mice (n=7) were immunostained for glucagon, insulin and Rab11A. **(E)** Colocalization of glucagon and Rab11A. The Rab11A image was pseudocoloured red for visualization of colocalization in the merged image and inset. **(F)** Colocalization of insulin and Rab11A. The Rab11A image was pseudocoloured green for visualization of colocalization in the merged image and inset. **(G)** Islets from non-diabetic mice were immunostained for glucagon, insulin and Rab11B. **(H)** Colocalization of glucagon and Rab11B. The Rab11B image was pseudocoloured red for visualization of colocalization in the merged image and inset. **(I)** Colocalization of insulin and Rab11B. The Rab11B image was pseudocoloured green for visualization of colocalization in the merged image and inset. **(J)** Islets from diabetic mice were immunostained for glucagon, insulin and Rab11B. **(K)** Colocalization of glucagon and Rab11B. The Rab11B image was pseudocoloured red for visualization of colocalization in the merged image and inset. **(L)** Colocalization of insulin and Rab11B. The Rab11B image was pseudocoloured green for visualization of colocalization in the merged image and inset. All images were acquired and post-processed as described in Methods. In each merged panel, selected areas (white squares) were magnified to show individual cells within islets. (**M**) Analysis of colocalization of glucagon and Rab11A or Rab11B, and Stmm2 and Rab11A or Rab11B in normal and diabetic (STZ) islets. Pearson’s correlation coefficient (PCC) values are shown as means ± SEM. Each dot represents a mean of 9-15 images per mouse.

### Depletion of lysosomes decreases glucagon secretion

To examine the underlying mechanisms of the endo-lysosomal system in glucagon trafficking in diabetes, we cultured αTC1-6 cells in media containing 16.7 mM glucose for 2 weeks, then treated them with Bafilomycin A1 (BFA1) for 2h to inhibit lysosomal activity. Unexpectedly, BFA1 markedly reduced the extent of K^+^-stimulated glucagon secretion (~ 6 times) (Figure 11).

**Figure 11:**
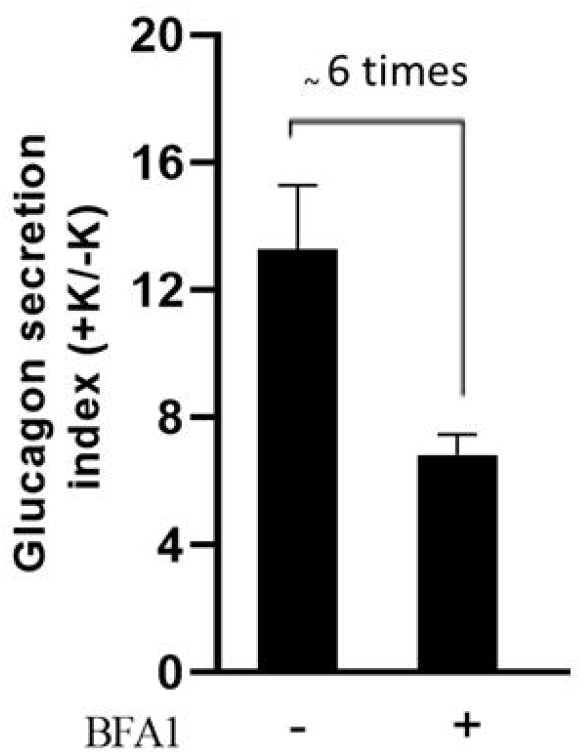
(A) Inhibition of lysosomes suppresses K^+^-stimulated glucagon secretion. αTC1-6 cells were cultured in high glucose (16.7 mM) condition for a long-term to mimic glucagon hypersecretion of diabetes. Cells were treated with medium (WT) or the lysosome inhibitor, Bafilomycin A1(10 nM, 2h; BFA1). Secreted glucagon was assayed by ELISA following 20 min incubation in the absence or presence of 55 mM KCl (baseline, and stimulated glucagon secretion, respectively). To calculate the glucagon secretion index, baseline secretion values were set to 1, and then corresponding fold changes in the stimulated conditions were calculated. Values were expressed as mean ± SD (n=4).

### Glucagon and Stmn2 co-localize to a Lamp1^+^ lysosomal compartment in diabetes mimicking αTC1-6 cells

To more fully explore the connection between lysosomal trafficking and glucagon hypersecretion in diabetes, we hypothesized that glucagon may be secreted through Lamp1^+^ lysosomes, which have been implicated in secretion of lysosomal cargo (Andrews N.W., 2017; Nakashima et al., 2019; Tseng et al., 2017). We therefore immunostained αTC1-6 cells for the lysosomal transmembrane protein, Lamp1, together with Stmn2 and glucagon (Figure 12A). Lamp1^+^ immunofluoresence was prominent at the cell membrane (Figure 12A), along with significant Stmn2 (Figure 12B) and glucagon (Figure 12C) immunofluorescence. Pearson’s correlation analysis showed a strong colocalization between Lamp1 and Stmn2 (Figure 12B; PCC 0.63 ± 0.04), and a high moderate colocalization between Lamp1 and glucagon (Figure 12C; PCC 0.50 ± 0.03).

**Figure 12:**
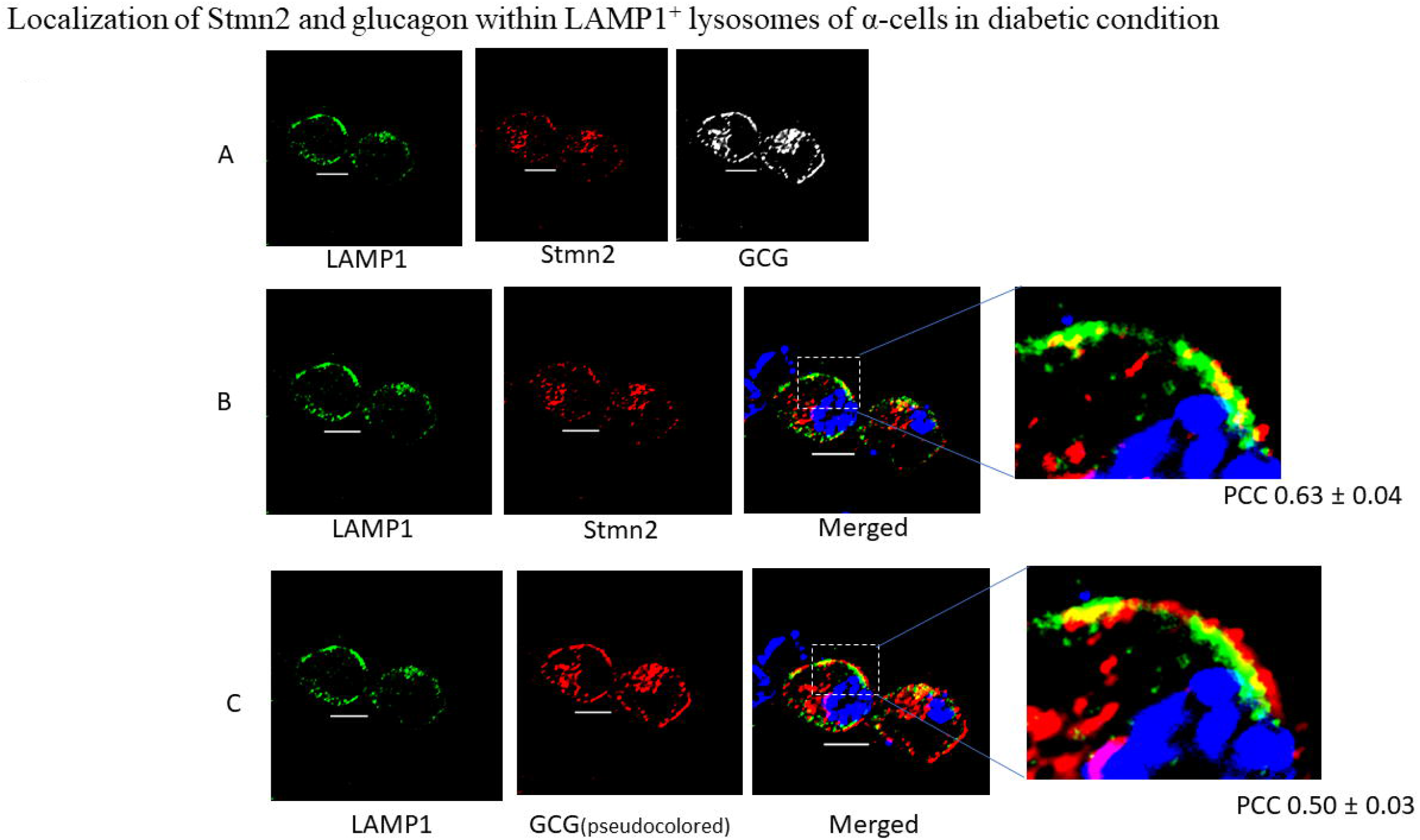
Glucagon and stathmin-2 are localized within Lamp1^+^ lysosomes in αTC1-6 cells in high glucose conditions. αTC1-6 cells were cultured in media containing 16.7 mM glucose for 2 weeks to mimic diabetes-induced glucagon hypersecretion. (A) Cells were immunostained with antibodies against glucagon (GCG), stathmin-2 (Stmn2), and lysosomal marker (Lamp1). (B) Colocalization of Stmn2 and Lamp1. (C) Colocalization of GCG and Lamp1. The GCG image was pseudocoloured red for visualization of colocalization in the merged image and inset. All images were acquired and post-processed as described in the Methods section. In each merged panel, a selected area (white square) was magnified to visualize the colocalization. Pearson’s correlation coefficient (PCC) value was shown as mean ± SEM.

## Discussion

Diabetes is always accompanied by a degree of hyperglucagonemia, which reflects dysregulated glucagon secretion from α-cells. One mechanism underlying defective glucagon secretion may be through impaired intracellular trafficking of glucagon. We have recently identified a novel protein in the alpha cell, stathmin-2 (Stmn2), that may be a negative regulator of glucagon secretion by directing glucagon to the endolysosomal system in αTC1-6 cells and islets from non-diabetic mice (Asadi and Dhanvantari, 2020). In the present study, we show that in diabetes, the abnormally high secretion of glucagon is accompanied by a relative decrease in cellular Stmn2. There was a concomitant and sharp reduction in the trafficking of both glucagon and Stmn2 into the Lamp2A^+^ lysosome, and increased localization of Stmn2 within the Rab7+ late endosome; however, neither protein was present in the early or recycling endosomal compartments. Interestingly, experiments in αTC1-6 cells grown in high-glucose media to mimic diabetes showed that inhibition of lysosomal biogenesis decreased glucagon secretion, leading us to search for another secretory compartment. We found that glucagon and Stmn2 localized in what may be secretory lysosomes, marked by Lamp1 immunoreactivity. We propose that, in diabetes, glucagon hypersecretion may result from a switch from trafficking to degradative lysosomes to secretory lysosomes, in addition to enhanced trafficking through secretory granules.

Our findings indicate that intracellular trafficking of glucagon through the endolysosomal system could be a new regulatory mechanism for glucagon secretion in pancreatic α-cells. In nondiabetic α-cells, glucagon secretion is regulated by several factors (Gromada et al., 2007): nutritional elements, such as glucose, amino acids and free fatty acids (Gromada et al., 2004; Gylfe and Gilon, 2014; Harp et al., 2016; Quesada et al., 2008; Ravier and Rutter, 2005; Salehi et al., 2006), neuronal effectors, such as norepinephrine and acetylcholine (Cryer, 2012; Quesada et al., 2008), and hormonal stimuli, particularly through paracrine regulation by insulin and somatostatin, and autocrine regulation by glucagon itself (Brereton et al., 2015; Briant et al., 2016; Quesada et al., 2008; Rodriguez-diaz et al., 2020; Walker et al., 2011; Watts et al., 2016). Diabetes disrupts this fine regulation of glucagon secretion, resulting in glucagon hypersecretion and aggravation of hyperglycemia. Also, alterations in the paracrine control of glucagon secretion by insulin results in an abnormal α-cell response to high glucose concentrations, either through insulin deficiency (Gilon, 2020; Lee et al., 2012) or α-cell insulin resistance (Gilon, 2020; Lee et al., 2014). Additionally, hepatic glucagon receptor resistance and impairment of amino acid turnover increases glucagon secretion from α-cells through the liver-α-cell axis (Adeva-andany et al., 2019; Janah et al., 2019). As well, higher levels of secreted glucagon and concomitantly co-secreted compounds such as glutamate from diabetic α-cells exacerbate glucagon hypersecretion in an autocrine manner (Gaisano et al., 2012).

The search for novel regulators of glucagon secretion has revealed that proteins associated with the α-cell secretory pathway may also play key roles in mechanisms that underlie hyperglucagonemia. Exposure of αTC1-6 cells to chronically high glucose concentrations resulted in an up-regulation of several secretory granule proteins, including processing enzymes, chromogranins and exocytotic proteins (McGirr et al., 2005), indicating that many components of the α-cell secretory pathway contribute to glucagon hypersecretion of diabetes. Our previous study on secretory granule proteomics in αTC1-6 cells revealed a dynamic profile of proteins predicted to associate with glucagon as possible mediators of glucagon secretion (Asadi and Dhanvantari, 2019), and our recent work has shown that Stmn2 may be one such novel regulator (Asadi and Dhanvantari, 2020). Knockdown of Stmn2 enhanced glucagon secretion from αTC1-6 cells, indicating that Stmn2 could be a negative regulator of glucagon secretion. Our present results demonstrating a reduction in Stmn2 cell content in islets from STZ-treated male mice concomitantly with glucagon hypersecretion are consistent with these results. A similar database search has shown that another granule protein, brefeldin A-inhibited guanine nucleotide exchange protein 3 or BIG3, also regulated glucagon secretion from mouse pancreatic islets, and its depletion *in vivo* resulted in glucagon hypersecretion (Li et al., 2015), possibly through promoting secretory granule biogenesis or maturation. Therefore, proteins within the α-cell secretory pathway are emerging as prominent regulators of glucagon secretion by mediating the intracellular trafficking of glucagon, and may explain dysregulated glucagon hypersecretion in diabetes.

In the present study, we observed that the relationship between glucagon and Stmn2 was disrupted, indicated by increased glucagon and decreased Stmn2 cell content. An altered balance between glucagon and Stmn2 has also been reported in islets from patients with diabetes; α-cell RNA sequencing analysis showed a higher *Gcg: Stmn2* gene expression ratio in islets from people with type 2 diabetes compared to healthy subjects (Lawlor et al., 2017). Proglucagon gene transcription, glucagon synthesis and secretion are all highly responsive to prevailing glucose concentrations (Dusaulcy et al., 2016; Huang et al., 2013; McGirr et al., 2005; Wewer Albrechtsen et al., 2017), reflecting the hyperglucagonemic state of diabetes. However, it appears that Stmn2 mRNA and protein levels may not be responsive to glucose concentrations. A BLAST search of the promoter region of the *Stmn2* gene (GeneBank: AH000817; mouse *Stmn2* complete cds) against the sequence of the glucose response element in the mouse glucagon receptor gene (Portois et al., 1999) (GeneBank: AF229079.1; mouse *Gcgr* complete cds), did not reveal any sequence homology. Thus, the enhanced secretion of Stmn2 in diabetes reflects an elevation in exocytotic activity in hyperglucagonemia (Huang et al., 2013), and not increases in mRNA or protein synthesis.

In islets from diabetic mice, the decrease in Stmn2 was accompanied by alterations in the trafficking of glucagon and Stmn2 through the late endosome-lysosome pathway, as indicated by the dramatic decrease in localization of glucagon and Stmn2 in Lamp2A^+^ lysosomes. These results are consistent with those from our previous study showing that siRNA-mediated depletion of Stmn2 sharply decreased the localization of glucagon in lysosomes and enhanced glucagon secretion. In the context of dynamic movements of cargos within the endolysosomal system, the late endosome cargos can be transported to the lysosome (anterograde) or the plasma membrane (retrograde). Therefore, we reasoned that diabetes-induced glucagon hypersecretion could occur through a switch from anterograde to retrograde transport. Retrograde transport can occur in two ways: *i*) to the early endosome, recycling endosome and then the plasma membrane, or *ii*) to the Golgi apparatus and then the secretory pathway (Hsu et al., 2012; Seto et al., 2002). However, the lack of glucagon and Stmn2 localization in early endosomes and recycling endosomes suggests that retrograde transport towards the early endosome-recycling endosome may not occur in α-cells. Therefore, glucagon hypersecretion in diabetes could be a result of enhanced retrograde trafficking of glucagon and Stmn2 from the late endosome towards the TGN, and then through the regulated secretory pathway.

Our data show that, in diabetic islets, glucagon trafficking to late endosomes is not altered, as the co-localization of glucagon and Rab7 was unchanged. The increase in Stathmin 2 in the late endosome, together with a reduction in Lamp2+ lysosomes, suggests a reduction in trafficking of glucagon and Stmn2 to the lysosome via the late endosome, and increased retention, at least of Stmn2, in Rab7+ late endosome. This apparent uncoupling of Stmn2 and glucagon suggests that Stmn2 is required for the trafficking of glucagon to Lamp2+ lysosomes in the normal regulation of glucagon secretion, and that, in diabetes, this step is impaired. Interestingly, siRNA-mediated depletion of Rab7 did not affect glucagon secretion in αTC1-6 cells (data not shown). These results are similar to those in pancreatic beta cells, where shRNA-mediated knockdown of Rab7 had no effect on insulin secretion (Zhou Y, Liu Z, Zhang S, Zhuang R, Liu H, Liu X, 2020). Instead, the routing of insulin to degradative lysosomes occurred through Rab7-interacting lysosomal protein (RILP) mediating the fusion of insulin secretory granules with lysosomes (Zhou Y, Liu Z, Zhang S, Zhuang R, Liu H, Liu X, 2020). Whether such a mechanism operates in alpha cells remains to be elucidated.

In order to further determine the role of lysosomal trafficking of glucagon in diabetes, we cultured αTC1-6 cells in media containing 16.7 mM glucose to induce a diabetic phenotype of glucagon hypersecretion. Cells were treated with the vacuolar H^+^ ATPase inhibitor bafilomycin A1 to inhibit lysosome biogenesis, as previously done in pancreatic beta cells and insulin-secreting cell lines (Pasquier et al., 2019). We expected to see no difference in glucagon secretion, as our findings in diabetic islets predicted that glucagon would not be trafficked to the lysosome. Instead, glucagon secretion was significantly decreased. This result could be interpreted in two ways: 1) bafilomycin A1 was inhibiting secretory granule biogenesis through inhibition of acidification; or 2) another acidification-dependent secretory compartment was being inhibited by the actions of bafilomycin A. It is unlikely that our regimen of BFA1 treatment results in inhibition of secretory granules, as previous reports demonstrate inhibition of granule acidification and biogenesis after 22h of treatment(Taupenot et al., 2005)), and we treated cells for only 2h, as previously described for inhibition of lysosomal biogenesis and fusion with autophagosomes (Goginashvili et al., 2015)(Pasquier et al., 2019).

The decrease in high glucose-induced glucagon secretion upon BafA1 treatment indicates that glucagon secretory granules are fusing with a lysosomal compartment that is secretion-competent. Since trafficking to the Lamp2A+ lysosome is inhibited in diabetes, it is possible that glucagon is trafficked to a secretory lysosome. We used Lamp1 to identify lysosomes that might have a secretory function, as Lamp1 is found on non-degradative lysosomes, and these lysosomes are often found at the plasma membrane (Cheng et al., 2018)(Yap et al., 2018). In diabetes-mimicking αTC1-6 cells, Lamp1+ lysosomes appear to be distributed at the plasma membrane, together with glucagon+ secretory granules. Co-localization of Lamp1 and glucagon at the plasma membrane suggests some degree of fusion of secretory granules with Lamp1^+^ lysosomes. Membrane-adjacent Lamp1^+^ vesicles contain the Ca^2+^-sensitive SNARE protein synaptotagmin 7 in MIN6 and INS1E cells (Monterrat et al., 2007), and may play a role as a Ca^2+^ sensor in lysosomal fusion and exocytosis. It is known that that the SNARE proteins VAMP7, SNAP23 and syntaxin 4 function in lysosomal fusion and exocytosis (Rao et al., 2004). Therefore, regulated exocytosis via lysosomes, in addition to enhanced secretory granule exocytosis, may contribute to the up-regulation of glucagon secretion in diabetes. It has been documented that mutant huntington is secreted via Lamp1^+^ lysosomes in a synaptotagmin 7-dependent manner (Trajkovic et al., 2017). Although we did not show such a mechanism in the present study, the role of secretory lysosomes in hyperglucagonemia will be the subject of future studies.

One limitation of our study is the identification of different endolysosomal compartments with only one marker. Mapping of endolysosomes in neurons has revealed that Lamp1 associates with a number of compartments in the endolysosomal pathway, some of which do not contain lysosomal degradative enzymes(Cheng et al., 2018), and that differentiation of endolysosomal compartments is possible only with using more than one immunofluorescent marker (Yap et al., 2018). Therefore, a more complete mapping of glucagon trafficking through the endolysosomal system in diabetes will require the use of more than one lysosomal marker.

Bafilomycin A1 is also a known inhibitor of autophagic flux, preventing the fusion of autophagosomes with lysosomes(Mauvezin and Neufeld, 2015)(Klionsky et al., 2008). The decrease in high glucose-induced glucagon secretion upon BafA1 treatment indicates a role for macroautophagy in the regulation of glucagon secretion. It is now well documented that impaired autophagy in pancreatic beta cells contributes to the progression of both type 1 and type 2 diabetes, as studied in mouse models of obesity and diabetes (Pasquier et al., 2019), proinsulin misfolding and ER stress-induced diabetes (Bachar-Wikstrom et al., 2013); and in islets from NOD mice and human islets (Muralidharan et al., 2021). Interestingly, impaired autophagy leads to the fusion of insulin secretory granules directly with lysosomes (Goginashvili et al., 2015). In contrast, impaired autophagy may be a mechanism in the abnormal up-regulation of glucagon secretion from alpha cells in diabetes. Further investigation is needed to probe the role of autophagy in alpha cell homeostasis.

In conclusion, our findings suggest a proof-of-concept model (Figure 13) in which diabetes suppresses trafficking of glucagon and Stmn2 from the late endosome towards the Lamp2A^+^ lysosomes. Consequently, there is retrograde trafficking of Stmn2 and glucagon towards the late endosome, and enhanced exocytosis of glucagon via Lamp1^+^ lysosomes. These findings suggest that altered lysosomal trafficking of glucagon may be considered as a potential mechanism for glucagon hypersecretion of diabetes.

**Figure 13:**
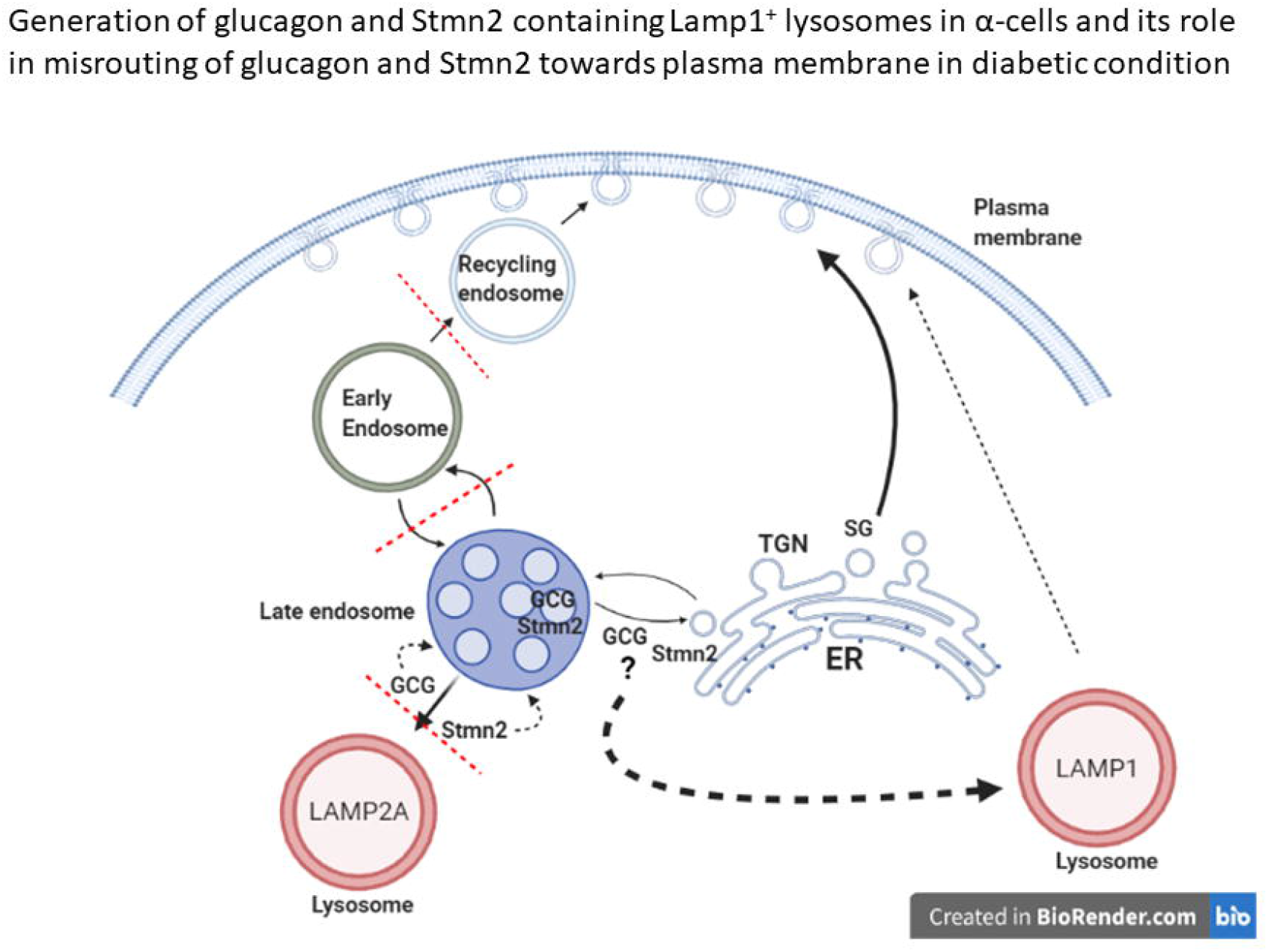
Enhanced glucagon secretion in diabetes results from a switch from trafficking to autophagic lysosomes to secretory lysosomes, in addition to enhanced trafficking through secretory granules. Glucagon (GCG) secretion is normally controlled by trafficking to the degradative Lamp2A^+^ lysosome via stathmin-2 (Stmn2). In diabetes, Stmn2 levels are reduced, and glucagon trafficking to Lamp2A^+^ lysosomes is inhibited. Glucagon secretion via secretory granules is enhanced, and in addition, glucagon is secreted through Lamp1^+^ lysosomes.

## Contribution statement

Designing and performing the experiments and analysis, writing the manuscript, preparing the figures and reviewing the manuscript prior to submission were done by FA and SD.

## Authors’ relationships and activities

The authors declare that there is no relationships or activities that may bring about bias, or results in bias for each part of the work.

## Acknowledgement

We would like to thank Ms. Karen Nygard and Mr. Reza Khazaee at Biotron Experimental Research Center, Western University for assistance with electron microscopy; Ms. Caroline O’Neil at Robarts Research Institute, Western University for tissue sample sectioning. We would like to acknowledge BioRender.com, which was used to generate the schematic figure.

